# A genetic approach to dissect the role of prefoldins in *Arabidopsis*

**DOI:** 10.1101/2021.01.28.428673

**Authors:** Noel Blanco-Touriñán, David Esteve-Bruna, Antonio Serrano-Mislata, Rosa María Esquinas, Francesca Resentini, Javier Forment, Cristian Carrasco-López, Claudio Novella-Rausell, Alberto Palacios, Pedro Carrasco, Julio Salinas, Miguel Á. Blázquez, David Alabadí

**Affiliations:** Instituto de Biología Molecular y Celular de Plantas (CSIC-Universidad Politécnica de Valencia), 46022 Valencia, Spain; Dipartimento di Bioscienze, Università degli Studi di Milano, 20133 Milano, Italy; Departamento de Biotecnología Microbiana y de Plantas, Centro de Investigaciones Biológicas (CSIC), 28040 Madrid, Spain; Departament de Bioquímica i Biologia Molecular, Universitat de València, 46100 Burjassot, Spain

**Keywords:** prefoldin complex, co-chaperone, TRiC/CCT, flowering time, stress, temperature, auxin

## Abstract

The prefoldin complex (PFDc) was identified in humans as co-chaperone of the cytosolic chaperonin TRiC/CCT. It is conserved in eukaryotes and is composed of subunits PFD1 to 6. PFDc-TRiC/CCT operates folding actin and tubulins. In addition to this function, PFDs participate in a wide range of cellular processes, both in the cytoplasm and in the nucleus, and their malfunction cause developmental alterations and disease in animals, and altered growth and environmental responses in yeast and plants. Genetic analyses in yeast indicate that not all functions performed by PFDs require the participation of the canonical complex. The lack of systematic genetic analyses in higher eukaryotes makes it difficult to discern whether PFDs participate in a particular process as canonical complex or in alternative configurations, *i.e*. as individual subunits or in other complexes. To tackle this question, and on the premise that the canonical complex cannot be formed if one subunit is missing, we have prepared an *Arabidopsis* mutant deficient in the six prefoldins, and compared various growth and environmental responses with those of the individual *pfd*. In this way, we demonstrate that the PFDc is required to delay flowering, for seed germination, or to respond to high salt stress, whereas two or more PFDs redundantly attenuate the response to osmotic stress. A coexpression analysis of differentially expressed genes in the sextuple mutant has identified several transcription factors, such as ABI5 or PIF4, acting downstream of PFDs. Furthermore, it has made possible to assign novel roles for PFDs, for instance, in the response to warm temperature.

## Introduction

Prefoldins (PFDs) are conserved proteins present in archaea and in eukaryotes that were identified in humans and in yeast as part of a hexameric complex, called PFD complex (PFDc; Vainberg *et al*., 1998; Geissler *et al*., 1998). PFDs can be classified into α- or β-type depending on their structure (Figure S1a) (Arranz *et al*., 2018). In eukaryotes there are two α-type (PFD3 and PFD5) and four β-type (PFD1, PFD2, PFD4, and PFD6) PFDs, whereas in archaea only one PFD per type is found. The PFDc adopts a jellyfish-like structure in which two α-subunits occupy a central position allowing the binding of four β-subunits (Siegert *et al*., 2000; Martin-Benito *et al*., 2002). In eukaryotes, the arrangement of the different subunits within the complex appears to be conserved (Gestaut *et al*., 2019).

Currently, the best characterized function of the PFDc is in proteostasis, as cochaperone of the chaperonin TRiC/CCT in the folding of tubulins and actin (Gestaut *et al*., 2019). Authors show that the substrate protein is transferred between the active sites of PFDc and TRiC/CCT until it is properly folded, avoiding the formation of deleterious protein aggregates. Yeast *gim/pfd* mutants show very similar cytoskeleton-related phenotypes, such as reduced a-tubulin levels (Vainberg *et al*., 1998; Geissler *et al*., 1998), which are not aggravated when several *gim/pfd* mutations are combined (Siegers *et al*., 1999). Tubulin-and actin-related phenotypes are also observed in *pfd* mutants in other model organisms. A missense mutation in the *PFDN5* gene causes developmental alterations in the central nervous system in mice, which are associated to reduced accumulation of α-tubulin and β-actin (Lee *et al*., 2011). Hypomorphic alleles of the *Drosophila MGR* locus, which encodes PFD3, cause defects in the formation of the meiotic spindle due to reduced tubulin levels, being this reduction also observed in fly DMEL-2 cells after knocking down *PFD4* (Delgehyr *et al*., 2012). Knock down of *PFD* genes in *C. elegans*, except *PFD4* that is divergent in this species, causes impaired cell division and embryo lethality due to defects in the rate of microtubule polymerization (Lundin *et al*., 2008). In *Arabidopsis, pfd* mutations provoke defects in the arrangement of cortical microtubules (MT) and in the formation of the phragmoplast, leading to impaired cell elongation and division, respectively (Gu *et al*., 2008; Rodriguez-Milla and Salinas, 2009; Perea-Resa *et al*., 2017). In summary, the similar phenotypes caused by mutations in individual *PFD* genes is consistent with the idea that the function of the PFDc is impaired when a subunit is missing. This view is further supported by the unique arrangement of subunits within the complex, based on specific protein-protein interactions (Gestaut *et al*., 2019).

Nevertheless, genetic analyses in yeast have shown that other functions of PFDs are not performed by the canonical PFDc. All PFDs, except GIM2/PFD3 and GIM4/PFD2, are required for transcription elongation of long genes and bind chromatin in a transcription-dependent manner (Millan-Zambrano *et al*., 2013). Furthermore, no additivity was found when combining affected *gim/pfd* mutants, which suggested that these PFDs may exert this role by being part of an alternative complex. In the same line, only GIM2/PFD3, GIM3/PFD4, and GIM1/PFD6 proteins are required for the transcription of genes in response to osmotic or oxidative stress (Amorim *et al*., 2017).

The implication of PFDs in other cellular processes is well documented (Liang *et al*., 2020), however, the lack of systematic genetic analyses makes it difficult to discern whether a particular role is exerted by the canonical PFDc or by individual subunits. For example, PFDN5/MM-1 acts as bridge protein that recruits a corepressor complex to the c-Myc transcription factor (Satou *et al*., 2001). Although the participation of pFdN5/MM-1 as c-Myc partner is demonstrated genetically (Fujioka *et al*., 2001) and the experimental data suggest that PFDN5/MM-1 fulfills this function, it cannot be ruled out that this role is performed as part of the PFDc. In *Arabidopsis*, PFD4 promotes the proteasomal degradation of the transcription factor HY5, and accordingly, its levels are augmented in *pfd4* mutants (Perea-Resa *et al*., 2017). HY5 levels are also increased in *pfd3* and in *pfd5* mutants, indicating that these other subunits are also involved. However, the limited genetic analysis precludes clarification as to whether this is a role performed by the canonical complex.

PFDs participate in diverse cellular processes in eukaryotes and their impaired function leads to disease and developmental abnormalities in animals (Liang *et al*., 2020), and to altered growth and response to environmental cues in plants (Rodriguez-Milla and Salinas, 2009; Perea-Resa *et al*., 2017; Esteve-Bruna *et al*., 2020) and in yeast (Millan-Zambrano *et al*., 2013; Siegers *et al*., 1999; Amorim *et al*., 2017). To understand the roles of PFDs in cellular processes, one of the issues that we need to clarify is whether they act as a canonical PFDc or as individual subunits in each case. To address this question in *Arabidopsis*, and since the function of the complex is impaired if one subunit is missing, we have prepared a mutant defective in the six PFDs and compared its growth habit and its behavior under various stresses with those of the individual mutants. Furthermore, we have identified novel functions for PFDs based on a transcriptomic analysis of the sextuple mutant.

## Results

### The PFDc forms *in vivo*

We first investigated whether the *Arabidopsis* PFDs can adopt the structure of their orthologs in yeast and humans. The structure of the *Arabidopsis* PFDs could be modeled *in silico* based on the structure of their human orthologs (Figure S1a) and assembled to form the jellyfish-like complex (Gestaut *et al*., 2019) (see the top view in Figure 1a and the comparison of the two complexes in Figure S1b). The high similarity between complexes suggests that the PFDc would adopt the same 3D arrangement in *Arabidopsis* and in humans *in vivo*.

**Figure 1.**
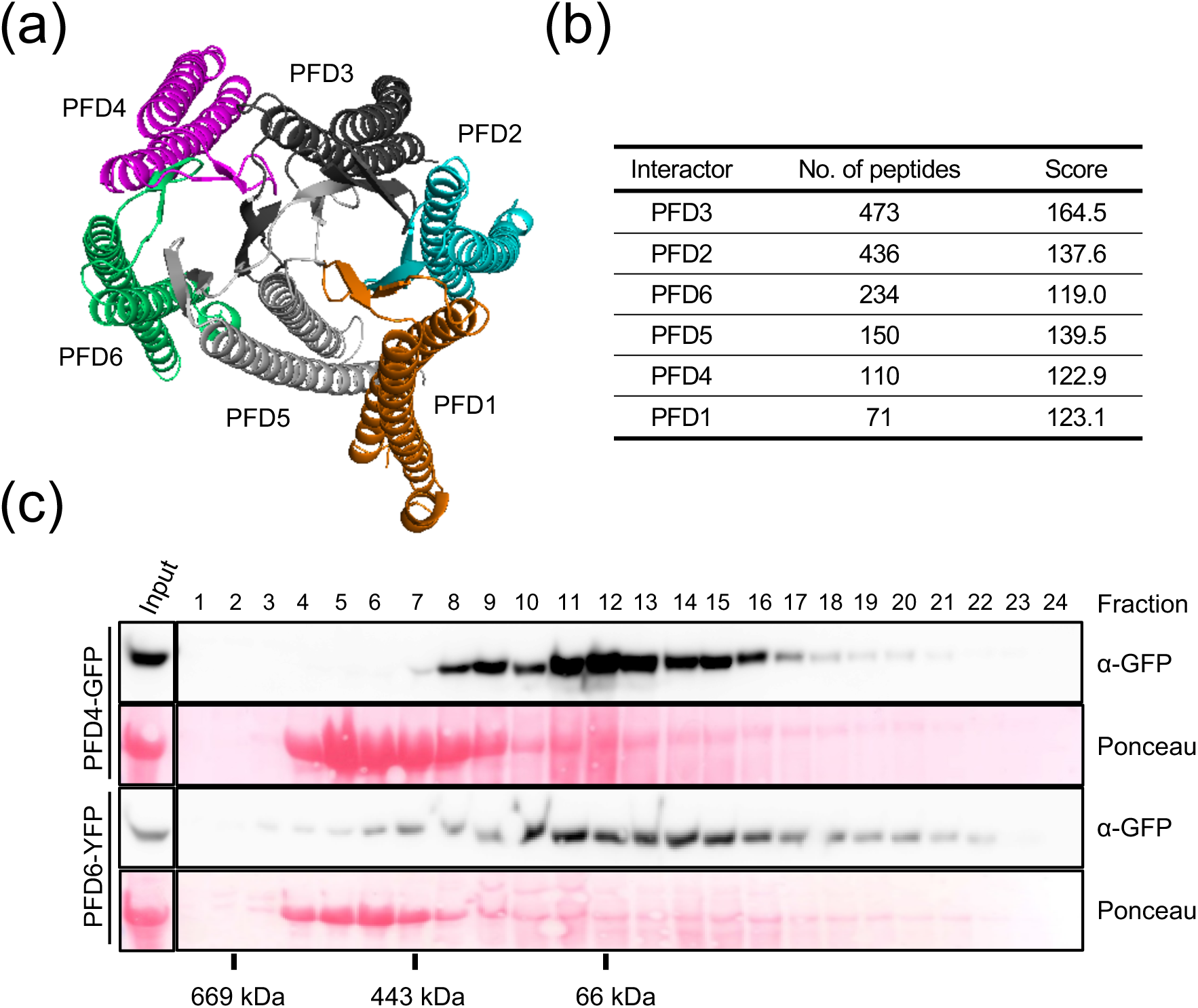
The *Arabidopsis* PFDc. (a) Predicted structure of the *Arabidopsis* PFDc using the human PFDc as template for modelling. (b) Identification of the *Arabidopsis* PFDc *in vivo*. The table summarizes the average number of peptides and the Mascot score corresponding to each PFD subunit after TAP of GS-PFD3. (c) Gel filtration fractions were analyzed by Western blot and the fusion proteins revealed with anti-GFP antibodies.

We next determined if the PFD proteins associate *in vivo*. For that purpose, we performed Tandem Affinity Purification using an *Arabidopsis* PSB-D cell suspension line expressing the GS-PFD3 fusion protein. After the two sequential immunopurification steps, the top PFD3 interactors identified in two replicates were the other five PFDs (Figure 1b), suggesting that the PFDc forms *in vivo*. Indeed, when we subjected extracts of seedlings expressing either *PFD4pro::PFD4-GFP* (Perea-Resa *et al*., 2017) or *35Spro::PFD6-YFP*(Esteve-Bruna *et al*., 2020) to gel filtration, the elution profiles of both proteins indicated that they may be incorporated into protein complexes of molecular weight compatible with the PFDc (ca. 130 KDa including the fusion protein) (Figure 1c).

### Preparation of the *6x pfd* mutant

In order to investigate the PFDs’ contribution to *Arabidopsis* development and response to the environment, and to determine in which cases PFDs participate as complex, we set out to prepare a sextuple mutant defective in the activity of the six PFDs. Mutants for *PFD* genes have been described in *Arabidopsis*, except for *PFD1* (Gu *et al*., 2008; Rodriguez-Milla and Salinas, 2009; Esteve-Bruna *et al*., 2020; Perea-Resa *et al*., 2017). We identified a T-DNA mutant for the *PFD1* gene in the GABI-Kat collection (Kleinboelting *et al*., 2012). The *pfd1* mutant carries the T-DNA inserted in the third exon (Figure S2a) and is null, or highly hypomorphic, as evidenced by the inability to amplify the full-length transcript by RT-PCR (Figure S2b). With available mutants for all *PFD* genes, we prepared the *pfd1 pfd2 pfd3 pfd4 pfd5 pfd6-1* sextuple mutant (hereafter referred to as *6x pfd*) by genetic crosses (see Methods section for details). An RNA-seq analysis of the sextuple mutant (see below) confirmed that *pfd1, pfd2, pfd3*, and *pfd4* alleles are null or highly hypomorphic and that *pfd6-1* carries the reported point mutation (Figure S3a). Nonetheless, it also showed that the *PFD5* gene was transcribed in the mutant, albeit at a reduced level (ca. 40% of wild type) (Figure S3a and S3b). This result contrasts with the absence of full-length *PFD5* transcript previously reported in the *pfd5* mutant (Rodriguez-Milla and Salinas, 2009). The insertion site is in the third intron (Figure S3c), suggesting that the T-DNA may be processed in a fraction of *PFD5* pre-mRNAs and that this occurs more often in the sextuple mutant than in the *pfd5*. After having obtained the *6x pfd* mutant, we investigated various growth and environmental responses in this mutant and compared them with those in individual *pfd*.

### PFDs participate in microtubule organization exclusively as canonical complex

Defects in the organization of MT have been described in *Arabidopsis* for the *pfd3, pfd4, pfd5*, and *pfd6* mutants (Gu *et al*., 2008; Rodriguez-Milla and Salinas, 2009; Perea-Resa *et al*., 2017). To determine if *pfd1* and *pfd2* mutations also cause defects in the MT organization, we introduced the microtubule marker *UBQ10::Venus-TUA6* (Salanenka *et al*., 2018) into both mutant backgrounds by genetic crosses. We imaged MT by confocal microscopy in two populations of cells in 3-day-old etiolated seedlings, apical hook cells, which have ceased elongation and have disorganized, randomly arranged MT, and in cells just below the apical hook, which undergo elongation and have organized MT, arranged parallel to the growth axis (Gu *et al*., 2008). MT were disorganized in apical hook cells in the wild type and organized in elongating cells below the hook (Figure 2a). The same MT organization was observed in the *pfd1* mutant, while MT were randomly arranged in both types of cells in *pfd2* seedlings. We analyzed other phenotypes dependent on tubulin folding: (i) the sensitivity to the microtubule-depolymerizing drug oryzalin and (ii) tubulin levels. *pfd1* seedlings showed similar response to oryzalin than the other *pfd* mutants, including *pfd2*, although reduced sensitivity was observed at the lowest concentration (Figure 2b). α-tubulin levels were reduced in all mutants (Figure 2c and d). In summary, these results show a similar behavior for all *pfd* mutants regarding microtubule-related phenotypes. The lack of apparent defects in the MT organization in the *pfd1* mutant may be due to the lower sensitivity of this assay to PFD1 deficiency compared to the other two.

**Figure 2.**
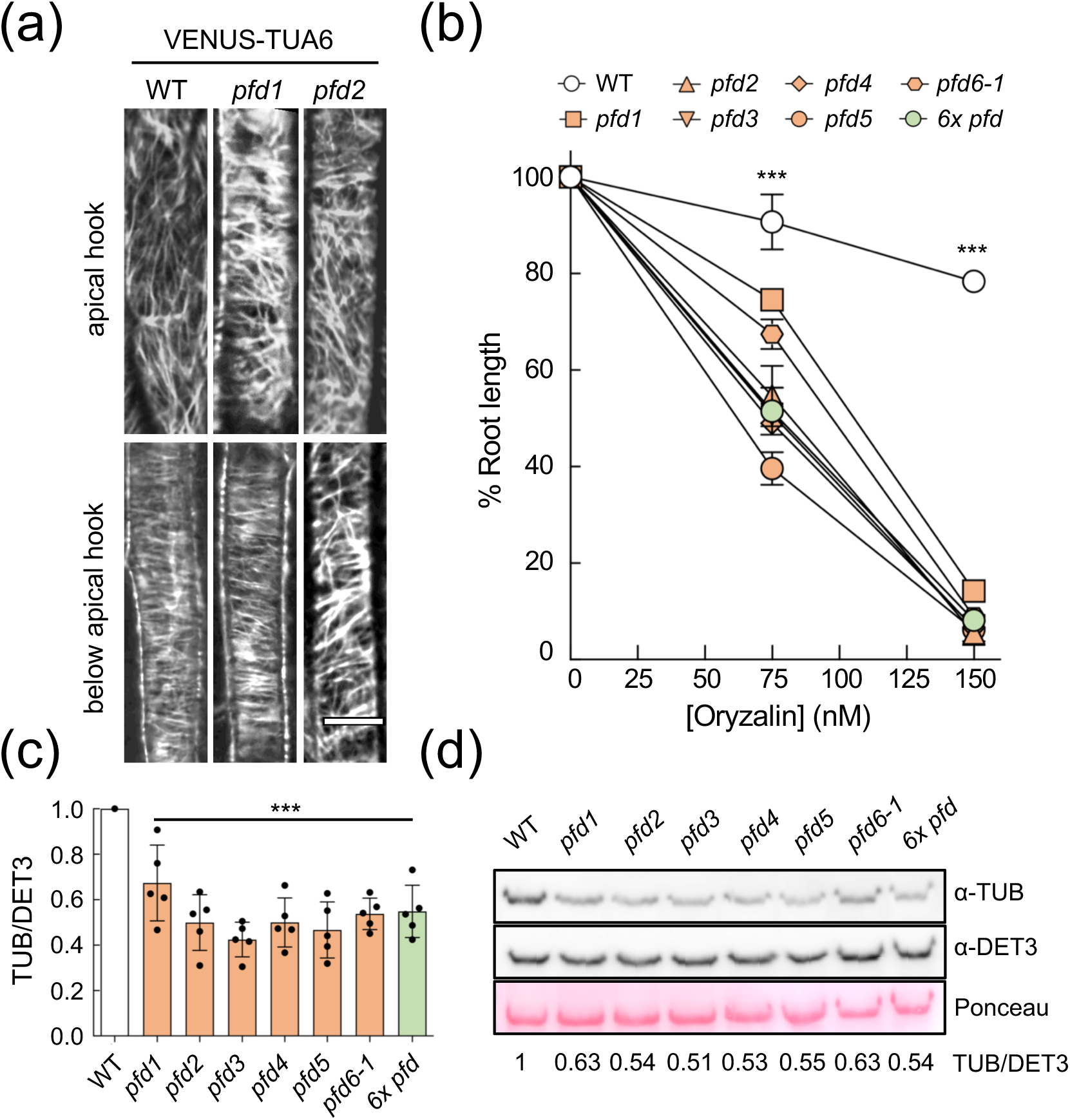
Microtubule alterations in *pfd* mutants. (a) Representative confocal images of VENUS-TUA6 in wild-type, *pfd1*, and *pfd2* hypocotyl cells. Scale bar = 10 μm. (b) Effect of oryzalin on root elongation. The graph shows the average of two biological replicates. Error bars indicate standard error of mean. (c) Representative Western blot showing α-tubulin levels in the wild type and in *pfd* mutants. DET3 was used as loading control. The ratio of tubulin/DET3 of this representative blot is shown. (d) Levels of α-tubulin relative to DET3. Data are the average of five independent experiments. Three asterisks represent P < 0.001 in Dunnet’s multiple comparison test after ANOVA test. *** indicates P < 0.001 in ANOVA tests when comparing all mutants with the WT.

We next investigated if this role is carried out by the PFDc. Studies in yeast and in humans indicate that it is involved (Gestaut *et al*., 2019). Nonetheless, the lack of genetic analyses makes it difficult to rule out a complex-independent role for the individual subunits. To tackle this question, we compared phenotypes of the *6x pfd* mutant with those of the individual mutants. On the premise that the complex cannot be formed if a subunit is missing, the rationale is that the microtubule-related phenotypes of individual *pfd* and the *6x pfd* mutants would be the same if the role of PFDs is performed entirely by the PFDc. The sextuple mutant showed the same sensitivity to oryzalin and a-tubulin levels than the individual *pfd* (Figure 1b-d). These results are the genetic demonstration that PFDs participate in microtubule-related processes exclusively as part of the PFDc.

### Complex-dependent and-independent contributions of PFDs to organ growth

Reduced growth is a common trait of *pfd* mutants (Perea-Resa *et al*., 2017; Rodriguez-Milla and Salinas, 2009; Esteve-Bruna *et al*., 2020; Gu *et al*., 2008). We next sought to determine if PFDs’ contribution to organ growth is mediated by the PFDc. We analyzed the size of the rosette and hypocotyl and root length in individual *pfd* mutants and in the *6x pfd*. The rosette size was reduced to a similar extent in individual *pfd* mutants and further reduced in the *6x pfd* (Figures 3a and b and S4). Microtubule-related defects leading to altered cell division and/or expansion may contribute to rosette growth alterations in individual *pfd* mutants. Nonetheless, the fact that the sextuple mutant exhibits a further reduction in size indicates that other processes, non-related to microtubules and controlled redundantly by two or more PFDs, are also altered in this mutant. PFDs therefore contribute to the rosette growth in two ways, dependent and independent of the PFDc.

**Figure 3.**
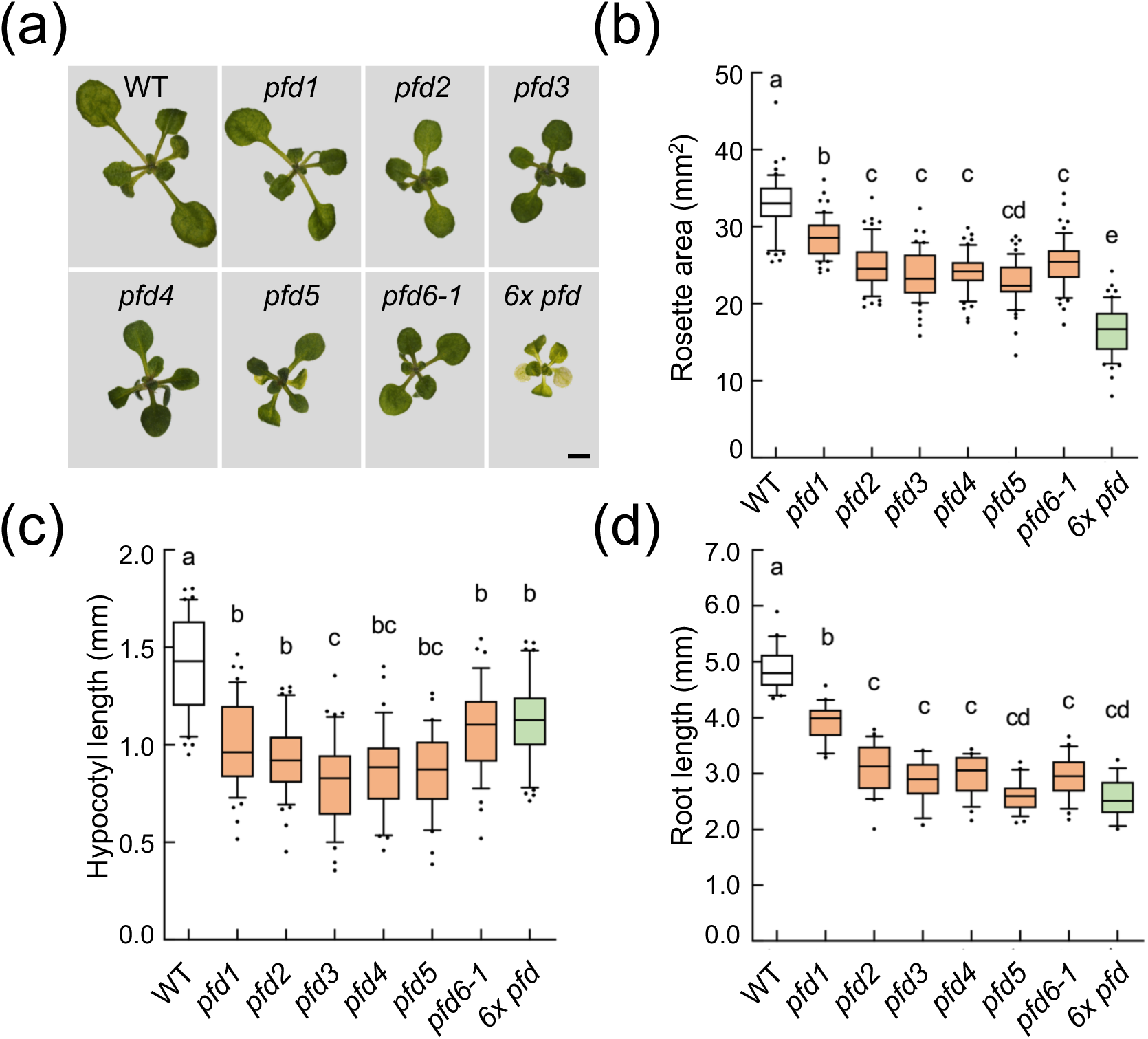
Effect of *pfd* mutations in organ growth. (a) Representative images of 14-day-old rosettes of the indicated genotypes grown in ½ MS plates under continuous light. Scale bar = 2 mm. (b-d) Box plots showing the rosette area (n ≥ 52) in 14-day-old plants (b), the hypocotyl length after growing 7-day-old etiolated seedlings (n ≥ 21) (c), and the root length (n ≥ 17) after growing 7 days in LD photoperiod (d). Horizontal lines inside boxes indicate the median. Whiskers indicate the highest and lowest values excluding outliers (points outside whiskers). Genotypes with different letters show significant differences at *P* < 0.05 according to ANOVA with Tukey’s HSD test.

The analysis of the hypocotyl and root length revealed a similar reduction in the size of both organs in individual *pfd* mutants and in the sextuple (Figure 3c and d). Despite the slight but significant differences among genotypes, the lack of an additive effect in the *6x pfd* mutant suggests that PFDs act as complex to promote both hypocotyl and root growth. At least part of the contribution of the PFDc to hypocotyl elongation may be mediated by its role in MT organization, since the growth of this organ is almost entirely mediated by cell expansion (Gendreau *et al*., 1997). In addition to cell expansion, cell divisions occurring in the root meristem also contribute to the growth of this organ. Therefore, PFDs contribute to root growth as complex that may mediate, at least, microtubule-dependent cell division and cell elongation.

### The PFDc contributes to the regulation of flowering time

We reasoned that the increased expression of *PFD* genes in the vegetative rosette and in the shoot apex, before and after the transition to flowering, would be compatible with a role for PFDs in flowering time regulation (Figure S5) (Winter *et al*., 2007). To test this hypothesis and to determine eventually whether this role is performed by the PFDc, we measured the flowering time of all individual *pfd* mutants and of the sextuple grown in short days (SD). Results show that all mutants flowered earlier than the wild type (Figure 4a). The phenotype was similar for all mutant lines, albeit the effect of the *pfd1* mutation was milder. Importantly, the absence of additive effects in the *6x pfd* suggests that the activity of PFDs on flowering time is exerted by the PFDc. This effect is independent of the photoperiod, since early flowering was also observed when the *6x pfd* mutant was grown under long days (LD) (Figures 4b and S6a).

**Figure 4.**
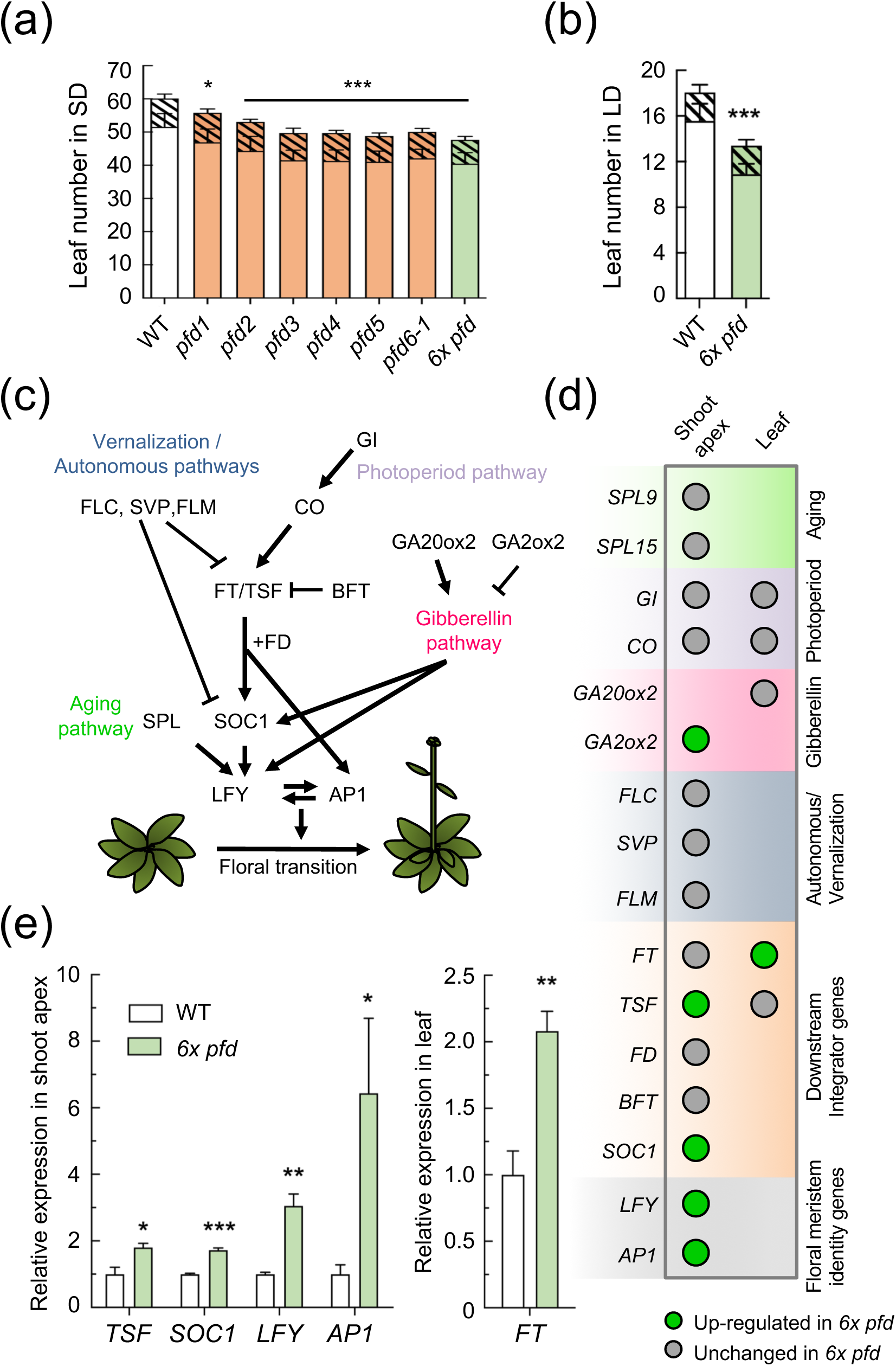
Flowering phenotype of *pfd* mutants. (a-b) Leaf number at bolting of plants grown in SD (n ≥ 9) (a) or in LD (n = 12) (b). Open and filled bars represent rosette and cauline leaves, respectively. Error bars indicate standard deviation of each kind of leaves. * and *** indicate *P* < 0.05 and 0.001, respectively, in Dunnet’s multiple comparison tests after ANOVA tests when the total number of leaves is considered. (c) Major pathways controlling flowering time in *Arabidopsis*. (d) Summary of qRT-PCR results. Circles indicate the genes analyzed in the apex shoot and /or the 2^nd^ oldest leaf of 14-d-old wild-type and *6x pfd* plants grown in LD. (e) Expression of misregulated genes in *6x pfd* plants compared to the wild type. Data are mean from 3 biological replicates. Error bars represent standard error from means. One, two, and three asterisks represent *P* < 0.05, 0.01, and 0.001 in t-tests.

Next, to try understanding how the PFDc contributes to the flowering time, we investigated whether the expression of key regulatory genes is altered in the *6x pfd* mutant (Figure 4c). The analysis included *SPL9* and *SPL15* (aging pathway); *FLM, SVP*, and *FLC* (vernalization and autonomous pathways); *GI* and *CO* (photoperiod pathway); *GA20ox2* and *GA2ox2* (gibberellin pathway); *FT, FD, TSF, BFT*, and *SOC1* (integrator genes); and *LFY*and *AP1* (meristem identity genes) (Fornara *et al*., 2010). We analyzed their expression by RT-qPCR in shoot apexes and/or in the second oldest rosette leaf from 14-day-old plants grown in LD. Among representative genes of different pathways, only the expression of *GA2ox2* was altered in the mutant (Figures 4d and S6b and c). We identified, nonetheless, the FT-TSF module as the main target of the PFDc. The expression of *FT* in leaves and of *TSF* in the shoot apex was higher in the sextuple mutant than in the wild type, which would explain the higher transcript levels of the downstream genes *SOC1, LFY*, and *AP1* that lead to the floral transition (Figures 4d and e and S6b and c). These results suggest that the early flowering of the *6x pfd* mutant is associated to increased FT-TSF activity. The PFDc, therefore, is required to delay flowering by repressing the expression of integrator genes.

### Individual PFDs attenuate the cold acclimation response

We next investigated the behavior of the *6x pfd* and individual *pfd* mutants when exposed to environmental challenges. PFD3, PFD4, and PFD5 attenuate the acclimation to low temperatures (Perea-Resa *et al*., 2017). The individual *pfd1* and *pfd2* mutants showed the same freezing tolerance as *pfd3, pfd4*, and *pfd5*, whereas a wild-type response was observed for *pfd6-1* (Figure 5a and b). The *6x pfd* plants showed a similar behavior than *pfd1* to *pfd5*. Taken together, our findings suggest that (i) PFD6 is not essential in controlling the adaptive response to low temperatures, and (ii) PFDs do not participate as canonical complex in this response. Nonetheless, since the *pfd6-1* allele carries a missense mutation that does not appear to interfere with the *in silico* assembly of the PFDc (Figure 5c), an alternative possibility is that PFDs participate in the low temperature response as canonical complex, and that the *pfd6-1* mutation does not affect the contribution of PFD6 to the function of the complex in this process, contrary to what occurs in others (Figures 2–4).

**Figure 5.**
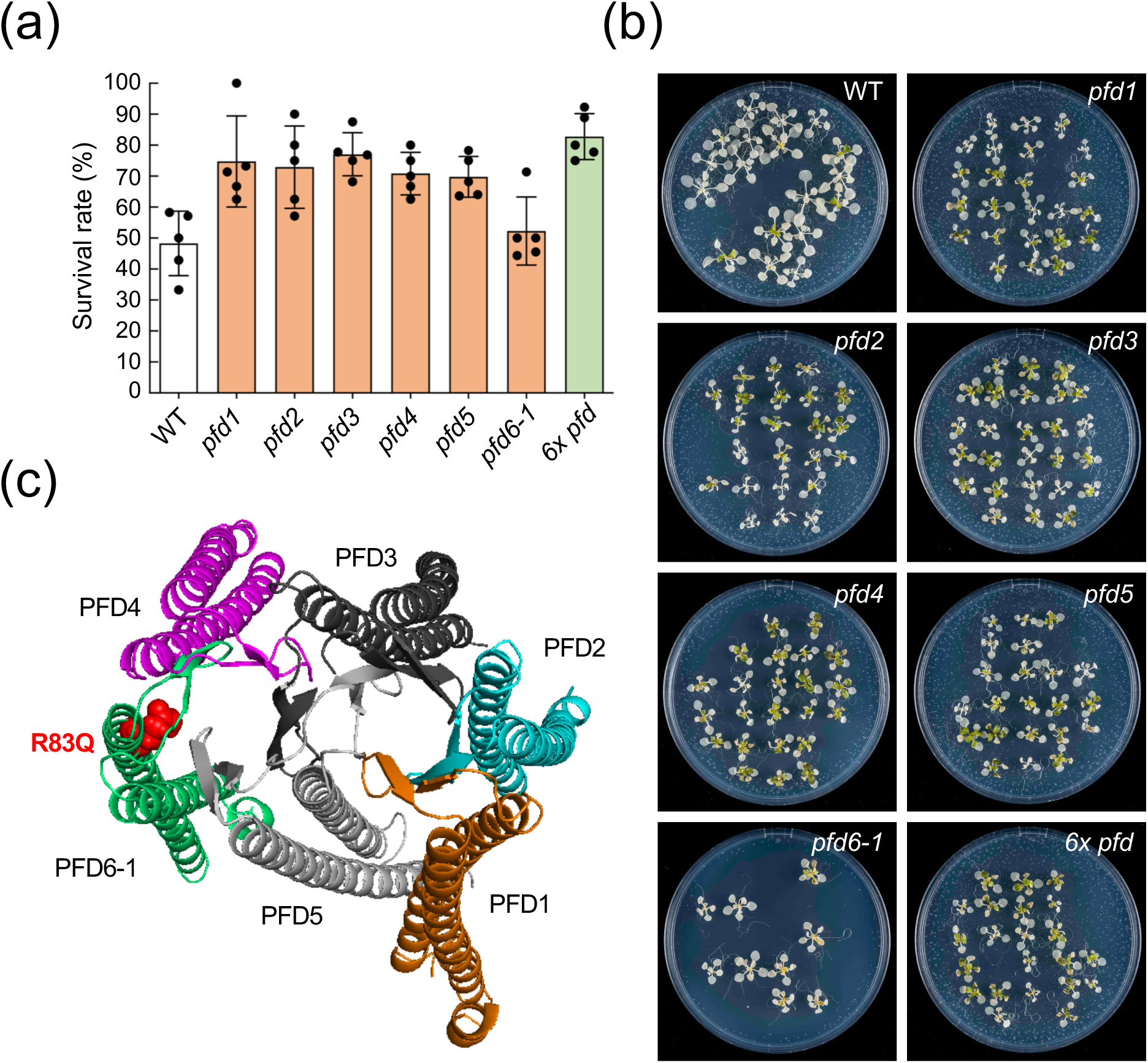
Cold acclimation in *pfd* mutants. (a) Freezing tolerance assay of cold-acclimated plants. Two-day-old wild-type, *pfd1, pfd2, pfd3, pfd4, pfd5, pfd6*, and *6x pfd* plants grown at 20°C were transferred to 4°C for 5 days and subsequently exposed to −12°C for 6 h. Survival rates were determined after 1 week of recovering at 20°C. Error bars indicate standard error of mean from five biological replicates. One, two, and three asterisks indicate *P* < 0.05, 0.01, and 0.001, respectively, in Dunnet’s multiple comparison tests after ANOVA tests. (b) Representative plates after recovery. (c) Three-dimensional reconstruction of the *Arabidopsis* PFDc using pfd6-1 instead of PFD6. The R-to-Q amino acid substitution in pfd6-1 is highlighted in red.

### Different contributions of PFDs to the response to salt and osmotic stress

The activity of PFD3, PFD4, and PFD5 is required for the plant’s response to salt stress (Rodriguez-Milla and Salinas, 2009; Esteve-Bruna *et al*., 2020). The root growth of *pfd2* and *pfd6-1* mutants was affected by the 100 mM NaCl treatment in a similar way to that of the *pfd3, pfd4*, or *pfd5* mutants, while it was less affected in the *pfd1* (Figure 6a). Interestingly, *6x pfd* seedlings behaved as the individual *pfd*, suggesting that PFDs contribute to the response to high salt as PFDc.

**Figure 6.**
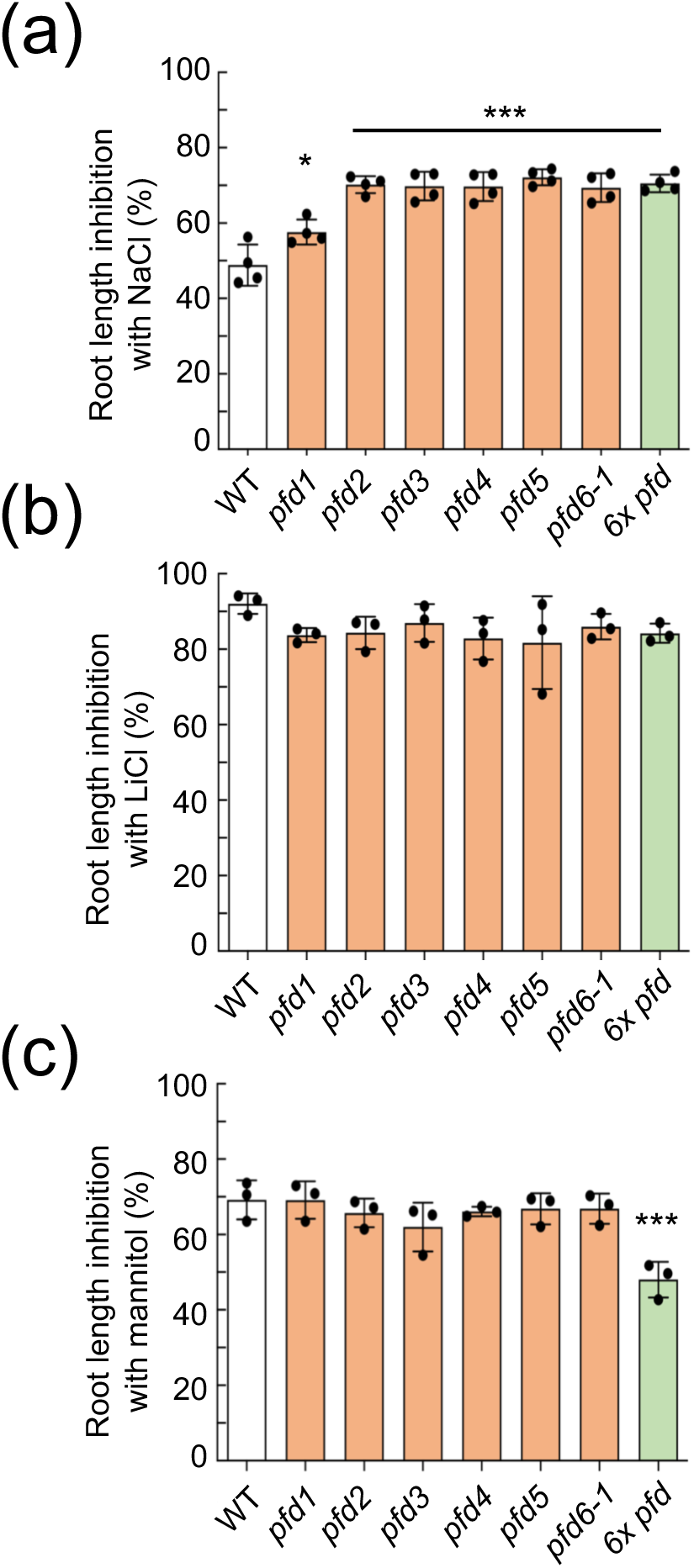
Response salt and osmotic stress in *pfd* mutants. (a-c) Root length in the presence of 100 mM NaCl (a), 12 mM LiCl (b), or (c) 300 mM mannitol. Bars represent mean from eight (a,c) or three (b) biological replicates. Error bars indicate standard error of mean. One and three asteriks indicate *P* < 0.01 and 0.001, respectively, in Dunnet’s multiple comparison tests after ANOVA.

The hypersensitive response to NaCl could be caused by either the ionic or the osmotic component of the treatment. Previous findings show that the sensitivity of *pfd3* and *pfd5* mutants is likely Na^+^-specific, because they are not hypersensitive to LiCl or mannitol (Rodriguez-Milla and Salinas, 2009). Nevertheless, it cannot be ruled out a general effect on ionic or osmotic stress if some PFDs act redundantly. We measured the root length of individual *pfd* and the *6x pfd* mutants in the presence of 12 mM LiCl or 300 mM mannitol. LiCl stress inhibited root growth in a similar way in all genotypes, ruling out any involvement of PFDs in the response to nonspecific ionic stress (Figure 6b). Interestingly, the *6x pfd* seedlings were more tolerant to the mannitol treatment, whereas a wild-type response was observed for individual *pfd* mutants (Figure 6c). These results indicate that the response of the plant to osmotic stress does not require the participation of the PFDc, but rather two or more PFD subunits that redundantly attenuate the response. Moreover, this result also indicates that it is unlikely that there is contribution of osmotic stress to the effect of high NaCl in *pfd* mutants.

The participation of PFDs in cold acclimation is reflected in augmented levels of *PFD4* transcript and protein in response to low temperatures (Perea-Resa *et al*., 2017). We investigated whether PFDs are subject to transcriptional or post-transcriptional regulation in response to salt or osmotic stress. Neither the expression of *PFD* genes nor levels of PFD4-GFP and PFD6-YFP proteins were affected by treatments with NaCl or mannitol, contrary to control genes or proteins (Figures S7 and S8).

### PFDs mostly contribute to gene expression independently of the PFDc

Impaired activity of PFDs results in altered gene expression (Esteve-Bruna *et al*., 2020). To determine whether PFDs participate in gene expression as complex, we compared the differential expressed genes (DEGs) identified by RNA-seq in the *pfd4* and *6x pfd* mutants. A total of 1186 DEGs, 734 up-and 452 down-regulated, were identified in the *6x pfd* mutant ([log_2_ FC] ≥ 2, *P* < 0.05; Figure 7a and b and Supplementary File 1). The relatively high number of misexpressed genes in the mutant highlights the relevance of PFDs’ activity for gene expression.

**Figure 7.**
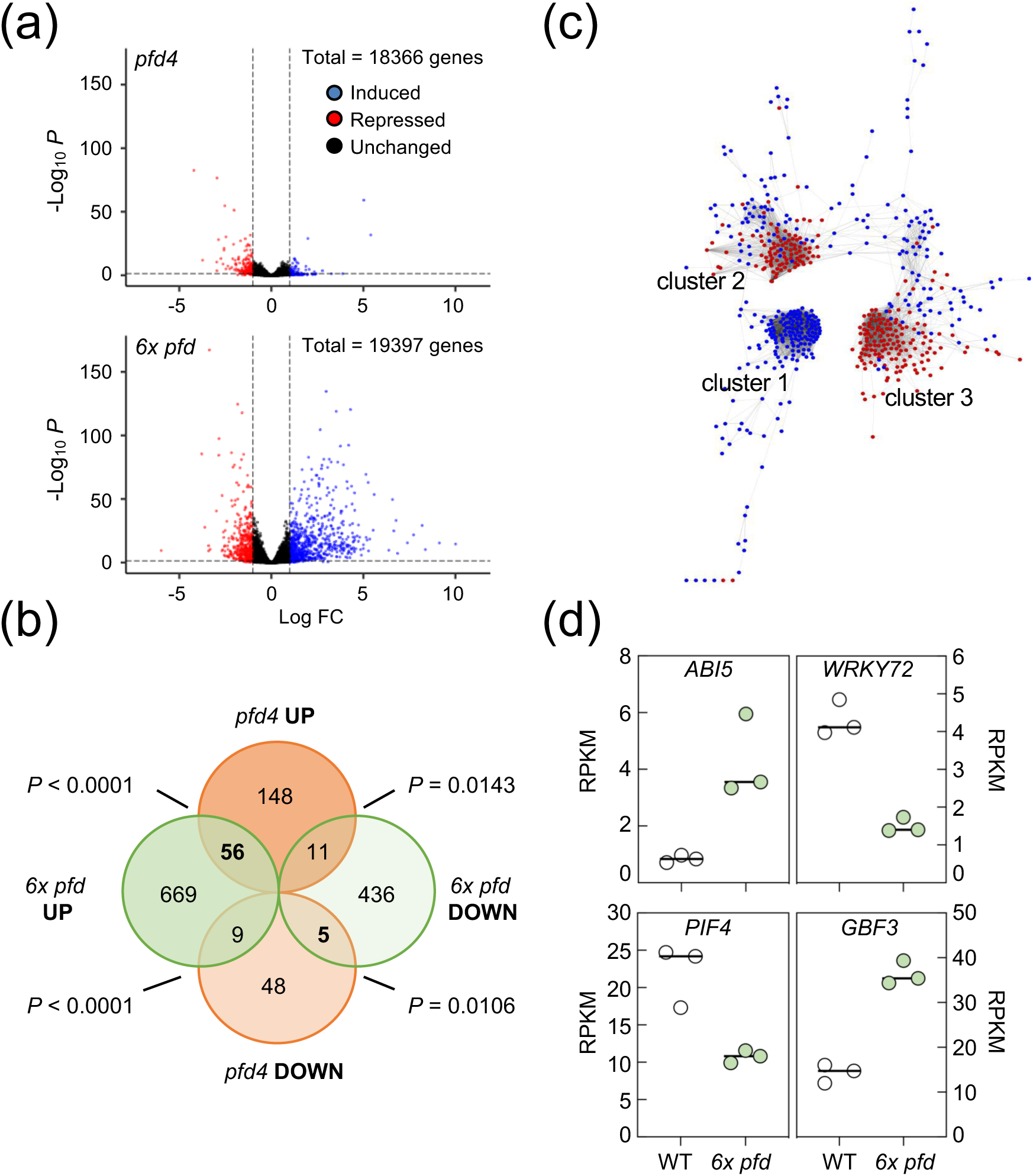
Transcriptomic analysis of the *6x pfd* mutant. (a) Volcano plots highlighting the DEGs in *6x pfd* and *pfd4* mutants (only genes with ≥ 1 RPKM in three replicates of mutants and/or wild type are shown). (b) Venn diagram comparing DEGs in *pfd4* and *6x pfd* mutants. (c) Cytoscape image of the coexpression network of DEGs in the *6x pfd* mutant. Red and blue indicates down-and upregulated genes, respectively. (d) Plots showing the expression level of the indicated genes extracted from the RNA-seq analysis. Each dot represents a replicate.

The number of DEGs was significantly lower in the *pfd4* mutant (198 DEGs, [log_2_ FC] ≥ 2, *P* < 0.05; Figure 7a and b and Supplementary File 2). A lower number of DEGs was also observed when the *pfd4* mutant was grown in soil instead of *in vitro* (117 DEGs, [log_2_ FC] ≥ 2, *P* < 0.05; Supplementary File 2; Esteve-Bruna *et al*., 2020). The fact that the molecular phenotype is more severe in the *6x pfd* mutant than in the *pfd4* suggests that the contribution of individual PFDs to gene expression is greater than that of the PFDc. It is important to note, however, that the majority of DEGs in *pfd4* seedlings, grown either *in vitro* or in soil, were not misexpressed in the *6x pfd* mutant (Figure 7b). Among the common genes, only the upregulated ones preferably followed the same trend (Figures 7b and S9a). The fact that the effect of the *pfd4* mutation on gene expression is less when found in a cellular context with mutations in all other *PFD* genes, suggests that part of the effect in the *pfd4* mutant may be due to overaccumulation of other PFDs.

### PFDs act upstream of a few TFs

We next wondered how PFDs affect gene expression. They may act via multiple pathways, but they may also act through few transcription factors (TFs) that control most of the PFD-dependent transcriptome. We reasoned that we could infer their behavior by looking at the architecture of the coexpression network of DEGs, which would likely organize itself into compact gene clusters if PFDs act through few TFs. We used the tool CORNET 3.0 (De Bodt *et al*., 2012) to determine the pairwise coexpression values among the 1186 DEGs in the *6x pfd* mutant in 454 microarray experiments. The analysis resulted in a network with 636 nodes and 9646 edges (Supplementary File 3), mostly organized into three compact gene clusters (Figure 7c). Interestingly, cluster 1 was exclusively formed by upregulated genes, while the two others mostly included downregulated genes (Supplementary File 4). The highest coexpression values were observed between genes of the same cluster (see Figure S9b for a larger image of clusters). This organization would be consistent with genes in each cluster being controlled by few TFs.

To investigate this possibility, we used the TF2Network tool that allows to identify putative regulatory TFs based in coexpression and DNA binding data (Kulkarni *et al*., 2018). We found that 80% of genes in cluster 1 are coexpressed with *ABI5* and 69% of them are direct targets of this TF. Importantly, *ABI5* itself was among the upregulated genes in the *6x pfd* mutant (Figure 7d and Supplementary File 1). ABI5 is a bZIP TF that plays a positive role in ABA signaling (Skubacz *et al*., 2016), and, in agreement with this, a GO analysis showed that the category “response to ABA stimulus” was overrepresented in cluster 1 (*P* = 1.5 × 10^-11^). Between 39% and 63% of downregulated genes of cluster 2 were coexpressed with *AtWRKY* TFs (*AtWRKY65, 36, 72, 35, 29, 9*, and *59*; cited from the most to the least coexpressed). *AtWRKY72* was also misexpressed in the mutant (Figure 7d and Supplementary File 1). The latter and AtWRKY29 have been characterized, being related to defense against pathogens (Bhattarai *et al*., 2010; Zhou *et al*., 2004). The analysis of cluster 3 showed poor coexpression values for any TF (less than 17%). Nonetheless, it revealed that 66% and 33% of genes were direct targets of GBF3 and PIF4, respectively. These TFs were also misexpressed in the *6x pfd* mutant (Figure 7d and Supplementary File 1). GBF3 is a bZIP TF that contributes to the plant’s response to abiotic stress (Ramegowda *et al*., 2017), and PIF4 is a well characterized bHLH TF that transmits information about ambient light and temperature to hormonal and growth pathways (Choi and Oh, 2016). In summary, the *in silico* analyses predicted that part of misregulated genes in the *6x pfd* mutant may be regulated by Ab15, WRKY72, GBF3, and PIF4.

### The transcriptome of the *6x pfd* mutant reveals novel functions for PFDs

To identify novel functions for PFDs, we searched for enriched Gene Ontology (GO) categories in the DEGs of *6x pfd* seedlings (Figure S9c, Supplementary File 1). The GO category “response to auxin stimulus” was more enriched in cluster 3 (*P* = 3.0 × 10^-9^) than when all DEGs were used (*P* = 2.1 x 10^-8^). Particularly striking is the presence of 13 auxin-responsive *SAUR* genes (Stortenbeker and Bemer, 2019) among those downregulated in cluster 3 (marked with an asterisk in Figure 8a). Indeed, another 13 *SAUR* together with the auxin biosynthesis gene *YUC8* and the auxin signaling gene *IAA29*, not represented in the microarray compendium used for coexpression analyses, also appeared downregulated in the *6x pfd* mutant (Figure 8a and Supplementary File 1). These results suggest that impairment of PFDs’ activity affects the auxin pathway.

**Figure 8.**
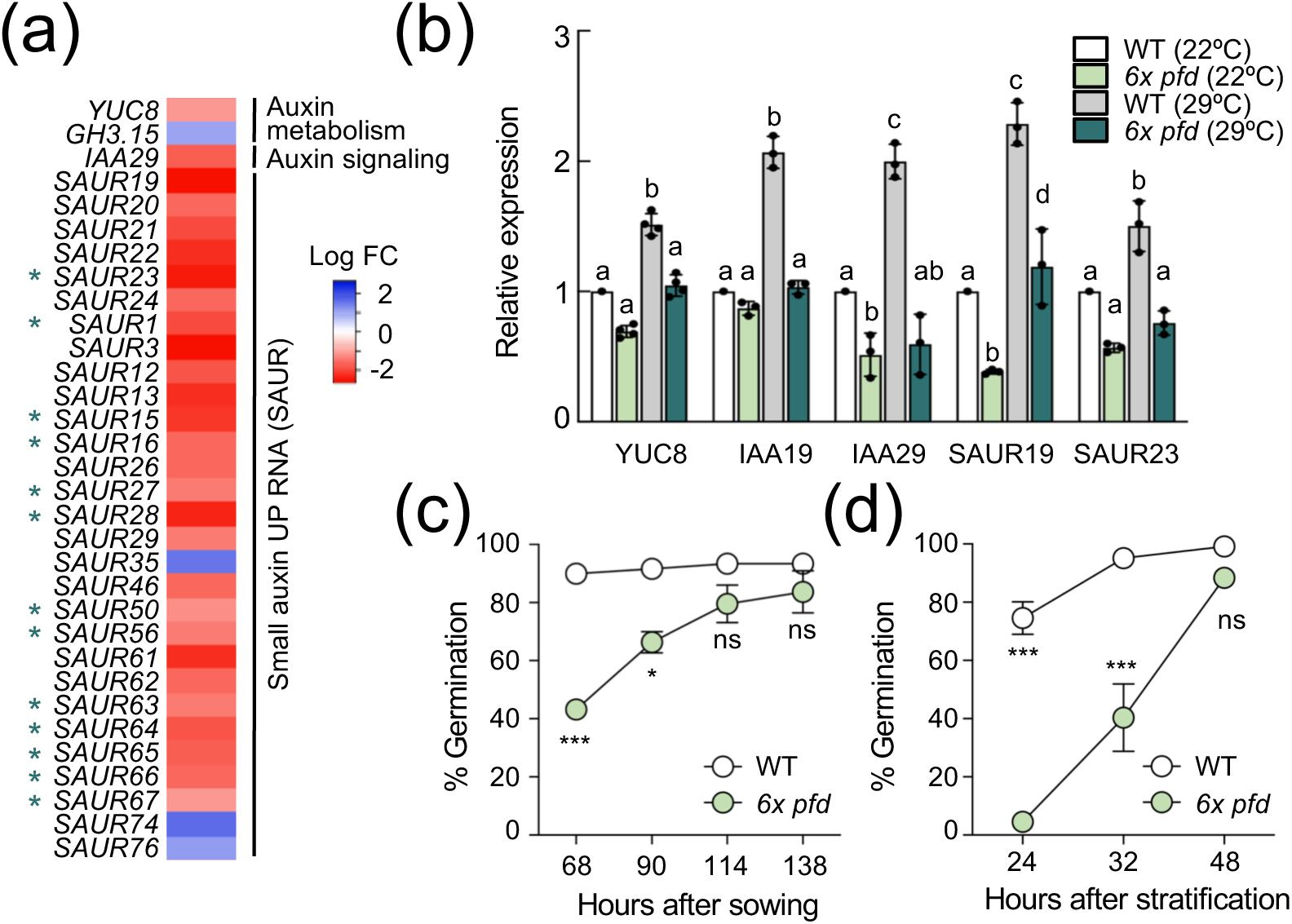
Novel functions for PFDs. (a) Heatmap showing the behavior of auxin related genes misregulated in *6x pfd* seedlings. Asterisks mark *SAUR* genes included in cluster 3. (b) Expression of auxin-related genes in response to 2 hours at 29°C. Data are mean of 3 biological replicates ± standard error of mean. Different letters show significant differences at *P* < 0.05 according to ANOVA with Newman-Keuls tests. (c, d) Germination rates of non-stratified (c) and stratified (d) wild-type and *6x pfd* seeds. Error bars represent standard error of mean from two biological replicates (each including 40-60 seeds). One, two, and three asterisks represent *P* < 0.05, 0.01, and 0.001 in Bonferroni tests after ANOVA tests, respectively. ns = no significant differences.

The TF2Network analysis revealed that PIF4 probably controls the expression of genes in cluster 3. PIF4 is important for the expression of auxin biosynthesis and signaling genes, especially in response to warm temperature (Quint *et al*., 2016). In agreement with this, the enrichment of the GO category “response to auxin stimulus” was most significant among the direct targets of PIF4 in this cluster (*P* = 1.2 × 10^-15^). The reduced expression of *PIF4* in the *6x pfd* mutant may contribute to the low expression of the auxin genes. Nonetheless, we reasoned that the consequences of low *PIF4* expression would be most obvious if we expose the *6x pfd* mutant to warm temperature. To test this, we selected five genes whose induction is PIF4-dependent: *YUC8* (Sun *et al*., 2012), *IAA19* (Huai *et al*., 2018), *IAA29* (Koini *et al*., 2009), and *SAUR19* and *23* (Franklin *et al*., 2011). *IAA19* was included in the analysis although its expression was not altered in the *6x pfd* mutant. The RT-qPCR analysis confirmed both the results of the RNA-seq and the thermal induction of genes (Figure 8b). Notably, the induction at 29°C was mostly impaired for all genes in the mutant, except for *SAUR19*. These results indicate that PFDs are required for the proper response of seedlings to warm temperature. The fact that *SAUR19* still responds to the temperature shift, despite it is dependent on PIF4 (Franklin *et al*., 2011), suggests that PFDs may affect the temperature response through other pathways in addition to those involving this transcription factor. The contribution of PFDs to the temperature response is not mediated by transcriptional regulation of *PFD* genes (Figure S10).

Since all *PFD* genes are transcriptionally active in imbibed seeds (Figure S5) and GO terms related to seed dormancy and germination were enriched in the DEGs in the *6x pfd* mutant (Figure S9c and Supplementary File 1), we wondered whether PFDs have a role in these processes. To determine the degree of dormancy and the germination capacity of *6x pfd* seeds, we compared their germination rate with that of the wild type with or without stratification at 4°C for 72 h. Seeds of both genotypes harvested at the same time were used. The *6x pfd* seeds exhibited an enhanced degree of dormancy compared with the wild type (Figure 8c). Furthermore, mutant seeds showed a delay in germination after stratification (Figure 8d). The role of PFDs in germination might depend on the PFDc because seeds of all individual *pfd* mutants showed a delay in germination similar to that of *6x pfd* seeds (Figure S11).

### The *6x pfd* mutant mimics transcriptional changes of wild-type plants exposed to stress

The enrichment of GO categories related to abiotic stress among DEGs in the *6x pfd* mutant (Figure S9c and Supplementary File 1) suggests that it may constitutively manifest transcriptomic stress features. To investigate this possibility, we compared the transcriptome of *6x pfd* seedlings with that of wild-type seedlings exposed to either 150 mM NaCl or 4°C for 24 h (Esteve-Bruna *et al*., 2020). The metaanalysis revealed that 56.7% of the DEGs in the mutant were altered in the wild type exposed to either of the two stresses (Figures 9a and Supplementary File 5), and that many of them behaved similarly (Figures 9b and S12). GO categories over-represented in the 163 DEGs common to the three conditions included response to abiotic stimuli and response to hormones (Figure 9c and Supplementary File 5). This indicates that PFDs are required to maintain adequate expression levels of many stress genes under non-stressful conditions.

**Figure 9.**
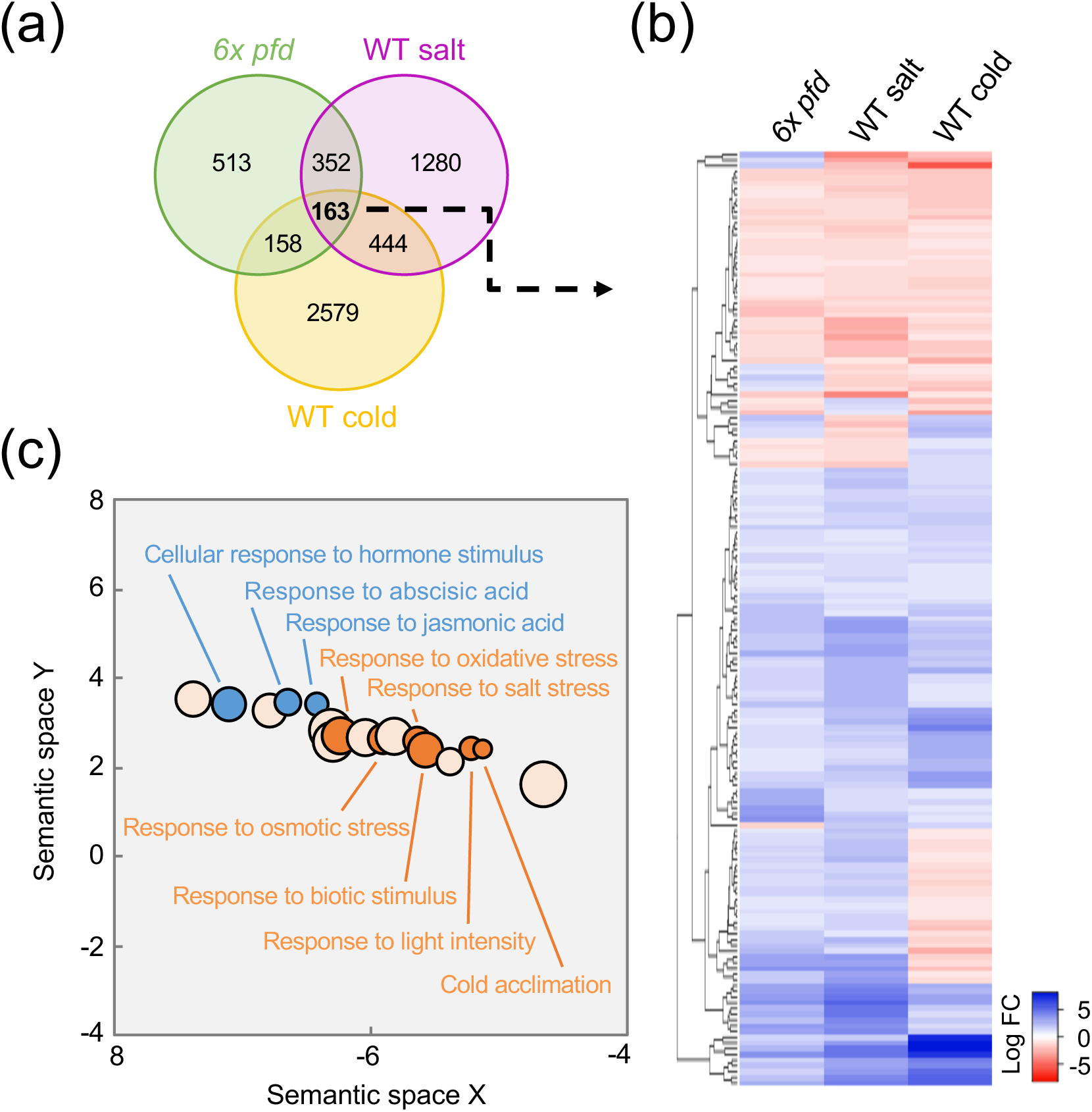
Constitutive stress signature in *6x pfd* seedlings. (a) Venn diagram comparing DEGs in the *6x pfd* mutant and in the wild type exposed to either salt or cold stress. (b) Heatmap showing the behavior of common DEGs in the three conditions. (c) GO terms enriched in the 163 common DEGs. Only some GO terms are represented. All GO terms are shown in Table S4. Bubble size is proporcional to the *P* significance of the GO enrichment.

Next, we hypothesized that PFDs are also required to achieve correct levels of gene expression in response to stress. To test this, we investigated the transcriptomic response of *pfd4* mutant seedlings exposed to either 150 mM NaCl or 4°C for 24 h. We found 2767 and 3718 DEGs in the *pfd4* mutant after salt and the low temperature treatments, respectively (Figure S13 and Supplementary File 6). Although many genes were common to the wild type, a clear *pfd4* signature was observed, since 1096 and 1121 DEGs were exclusively misexpressed in the mutant. These results indicate that at least PFD4 is required to reach proper expression level of many genes after the plant is subjected to stress.

## Discussion

PFDs are required for normal development of animals and their malfunction, due to mutation or misexpression, is normally associated to cancer or disease (Liang *et al*., 2020) or causes embryo lethality (Lundin *et al*., 2008; Delgehyr *et al*., 2012). *Arabidopsis pfd* mutants, including the *6x pfd*, are viable and do not show apparent developmental defects beyond reduced size, indicating that PFDs’ activity is mostly dispensable for normal development in this species (this work; Esteve-Bruna *et al*., 2020; Rodriguez-Milla and Salinas, 2009; Perea-Resa *et al*., 2017; Gu *et al*., 2008). Rather, their role in plants appears to be more relevant for properly interpreting environmental challenges (Esteve-Bruna *et al*., 2020; Rodriguez-Milla and Salinas, 2009; Perea-Resa *et al*., 2017). In fact, we show here that plants defective in the six PFDs perform better than the wild type when exposed to osmotic stress, which adds to known defects in the response to high salt or low temperature. It is important to note that our results expand this view. We show that they are not only required to respond to stress, but also to respond to environmental changes that in principle do not represent a stressful scenario for the plant, such as a moderate increase in the ambient temperature. This is further supported by the finding of enriched GOs related to environmental and stress responses in the gene set misregulated in the *6x pfd* mutant, and with the fact that the TFs identified by coexpression analyses acting downstream of PFDs mainly participate in environmental responses. The different effect of PFDs’ malfunction on the plant’s response to environmental changes, *i.e*. hypo-or hypersensitivity, is probably an indication of the varied modes of action through which these proteins act, which are just beginning to glimpse.

Determining whether PFDs participate in a particular process as members of the canonical PFDc or, conversely, in alternative configurations, *i.e*. other complexes or as individual subunits, provides clues to their mode of action. The analysis of the *6x pfd* mutant has allowed us to identify processes that are dependent and independent of the activity of the canonical PFDc. Joint action with the TRiC/CCT would be expected for the PFDc in those processes that require the participation of the canonical complex (Gestaut *et al*., 2019). We have found that several growth-involving processes depend on it, in which it probably participates through the folding of at least tubulins. For other processes also depending on the PFDc, such as the plant’s response to salt stress or the regulation of flowering time, the mechanistic connection with tubulins is not so obvious. Interestingly, altering microtubule polymerization affects gene expression in plants (Sangwan *et al*., 2001), and actin, another bona fide substrate of PFDc-TRiC/CCT, performs roles in the nucleus related to transcription (Blettinger *et al*., 2004). This opens up the possibility that the effect of the PFDc on the expression of the flowering integrator genes or of genes involved in salt stress is mediated through these two protein substrates. Alternatively, the PFDc may act through actin-related proteins, some of which are subunits of chromatin remodeling complexes that regulate the expression of *FT* (Kumar *et al*., 2012; March-Diaz and Reyes, 2009) and also of stress genes (Wang *et al*., 2019). In fact, an actin-related protein, in this case belonging to a cytosolic complex, is substrate for TRiC/CCT chaperonin in vertebrates (Melki *et al*., 1993).

PFDc-TRiC/CCT may fold substrates other than tubulins or actin that could also mediate the function of the PFDc. Although not supported by functional or genetic analyses, interactomic approaches suggest that PFDc-TRiC/CCT assist the folding of the histone deacetylase HDAC1 in the nucleus of human cells (Banks *et al*., 2018). Similar approaches have identified the PFDc and TRiC/CCT complexes associated to the TOR kinase complex in *Arabidopsis* (Van Leene *et al*., 2019). In *Drosophila*, TRiC/CCT interacts physically with members of the TOR complex and is required for TOR function, probably by participating in its assembly (Kim and Choi, 2019). That the TOR complex is substrate of the PFDc-TRiC/CCT in plants is an interesting possibility that awaits investigation.

We have identified other processes in which the action of PFDs does not involve the canonical complex. Genetic analyses indicate that two or more PFDs act redundantly to promote rosette growth or to attenuate the response to osmotic stress. In these cases, it is more difficult to anticipate their mode of action. They may participate as individual subunits or as part of alternative complexes. For example, our previous results show that PFDs can participate in proteostasis independently of the PFDc. PFD4 acts as adaptor to mediate the stabilization of the spliceosome complex LSM2-8 by the chaperone Hsp90, a process in which PFD2 does not appear to be involved (Esteve-Bruna *et al*., 2020). Their participation in alternative complexes is a very exciting possibility that has been proposed in yeast to explain the role of PFDs in transcriptional elongation, although the identity of the complex is currently unknown (Millan-Zambrano *et al*., 2013). A PFD-like complex formed by PFD2 and PFD6 together with the PFD-like proteins URI1, UXT, PDRG1, and ASDURF has been identified in animals (Chaves-Perez *et al*., 2018). It is therefore reasonable to think that the putative *Arabidopsis* PFD-like complex may participate in PFDc-independent processes requiring PFD2 and/or PFD6. Defining the *in vivo* interactome of PFDs in different *pfd* mutant backgrounds would allow identifying putative alternative complexes and their partners, helping therefore to delineate PFDc-independent mechanisms for PFDs’ action.

In summary, our results place PFDs as relevant players in the plant’s response to environmental changes. Furthermore, the genetic analyses provide evidence that PFDs’ action is not always mediated by the canonical PFDc, and clues about their possible mode of action in each case. The genetic resources generated in this work will help deciphering the mechanisms through which these versatile proteins act.

## Materials and methods

### Plant material and growth conditions

*Arabidopsis thaliana* accession Columbia-0 (Col-0) was used as wild type. Most mutants and transgenic lines have been described: *pfd2* (Esteve-Bruna *et al*., 2020), *pfd3* and *pfd5* (Rodriguez-Milla and Salinas, 2009), *pfd4* (Perea-Resa *et al*., 2017), *pfd6-1* (Gu *et al*., 2008), *RGApro::GFP-RGA* (Silverstone *et al*., 2001), *PFD4pro::PFD4-GFP* (Perea-Resa *et al*., 2017), *35Spro::PFD6-YFP* (Esteve-Bruna *et al*., 2020), and *UBQ10pro::VENUS-TUA6* (Salanenka *et al*., 2018). The *pfd1* mutant (GK-689A09) was obtained from the Nottingham Arabidopsis Stock Centre.

To grow seedlings *in vitro*, seeds were surface-sterilized, sown on plates with half-strength MS (Duchefa) media, pH 5.7, that includes 1% (w/v) sucrose and 8 g L^-1^ agar (control media), and stratified at 4°C for 3-4 days. Plates were exposed to continuous light (50-60 μmol m^-2^ s^-1^) or LD photoperiod (16 h of 90 μmol m^-2^ s^-1^) at 22°C. For hypocotyl length measurements and TUA6-VENUS visualization, seedlings were grown without sucrose.

To obtain the *6x pfd* mutant, all double mutant combinations were first prepared by crossing individual *pfd*. Then, double mutants were used to obtain five triple mutants (*pfd1,2,3; pfd1,3,5; pfd2,3,5; pfd2,4,6*, and *pfd3,5,6*). Triple mutants were used to obtain three quadruple mutants (*pfd1,2,3,5; pfd1,3,5,6*, and *pfd2,3,5,6*). Crosses between triple and quadruple mutants, and between quadruples, were performed to obtain three quintuples (*pfd1,2,3,5,6; pfd1,3,4,5,6*, and *pfd2,3,4,5,6*) and the sextuple mutant. All mutant combinations were genotyped in F2 generations with primers listed in Table S1.

The *UBQ10pro::VENUS-TUA6 pfd1* and *UBQ10 pro::VENUS-TUA6 pfd2* lines were obtained by crossing and confirmed by genotyping in F2 or F3 generations with the primers listed in Table S1.

### Protein Structure Prediction

The 3D structure of the *Arabidopsis* PFDc was modeled using the human PFDc (PDB code 6NR8) as template (Gestaut *et al*., 2019) using Modeller (release 9.23) (Webb and Sali, 2016), and visualized with PyMOL 2.4 software.

### Tandem affinity purification

The GS-PFD3 fusion under the control of the *35S* promoter was generated and used to transform *Arabidopsis* PSB-D cell suspension cultures. The sequential affinity purification was performed as described (Van Leene *et al*., 2015). Proteolysis and peptide isolation, acquisition of mass spectra, and protein identification was carried out at the Unidad de Proteómica (CNB, Madrid, Spain).

### Gel filtration and Western Blot Analysis

For gel filtration, protein extracts of 7-day-old *PFD4pro::PFD4-GFP* and *35Spro::PFD6-YFP* seedlings grown under continuous light were prepared in Extraction Buffer (50 mM Tris–HCl pH 7.5, 150 mM NaCl, 10 mM MgCl_2_, 10% glycerol, 0.5% Nonidet P-40, 2 mM PMSF and 1x protease-inhibitor cocktail) and loaded onto a Superose^™^ 6 Increase column (GE Healthcare). Twenty-four fractions of 0.5 mL were collected and precipitated as described (Esteve-Bruna *et al*., 2020). Proteins were separated in 12% SDS-PAGE and transferred to a PVDF membrane by Western blotting. Membranes were stained with Ponceau S solution and then incubated with anti-GFP antibody (JL-8 Takara Bio Clontech, Lot #A8034133).

To determine tubulin levels in *pfd* mutants, 7-day-old seedlings were grown under continuous light at 22°C. To determine GFP-RGA, PFD4-GFP, and PFD6-YFP levels in the presence of salt or mannitol, 7-day-old seedlings grown in continuous light were exposed to 100 mM NaCl or 300 mM mannitol in liquid media for 0, 8, or 24 h. Ground frozen tissue from whole seedlings was homogenized in Extraction Buffer. Total proteins were separated in 12% SDS-PAGE, transferred to PVDF membranes by Western blotting and visualized with anti-GFP (JL-8 Takara Bio Clontech, Lot #A8034133, 1:5000), anti-α-tubulin (Invitrogen, Lot #TD2479055, 1:1000), or anti-DET3 (1:10000, provided by Prof. Dr. Karin Schumacher). Quantification of protein bands in Western blots was performed using FIJI (https://fiji.sc/).

### Confocal microscopy

To determine MT organization, seeds of *UBQ10pro::VENUS-TUA6* in wild type, *pfd1*, and *pfd2* backgrounds were germinated in MS media without sucrose and grown vertically in darkness for 3 days at 22°C. MT were visualized in cells at the apical hook and immediately below by using a Nikon Eclipse Ti2 inverted microscope, equipped with a Yokogawa Spinning Disk Field Scanning Confocal System (https://www.microscope.healthcare.nikon.com/en_EU/products/confocal-microscopes/csu-series/specifications). The objective used was the oil immersion CFI 60 x H Plan Apocr λ oil W.D. 0.13 mm N.A. 1.40. VENUS was excited by a 488 nm single mode optical fiber laser and the emission was collected at 525-550 nm. Images were collected with a Photometrics Prime BSI CMOS camera (https://www.photometrics.com/products/prime-family/primebsi) with an exposure time of 100 to 200 ms with a 1×1 binning (2048 x 2048 pixels). The NIS-Element AR (Nikon, Japan, http://www.nis-elements.com/) was used as platform to control microscope, laser, camera, and post-acquisition analyses. Images were denoised by using the Denoise.ai algorithm (https://denoise.laboratory-imaging.com/process) and then analyzed using FIJI software.

### Phenotypic analyses

The quantification of the sensitivity to oryzalin was carried out by measuring the length of the primary root of 7-day-old seedlings grown in LD on vertical plates supplemented with increasing concentrations of oryzalin (0, 75, and 150 nM). Root length was measured using FIJI.

To determine rosette area, seeds were sown on 140-mm diameter Petri dishes at low density (40 seed per plate) and grown under continuous light. Plates were photographed with a digital camera 14 days after germination. The measurement was obtained using the FIJI plug-in Rosette tracker from at least 3 biological replicates (20 seedlings each). Representative images were taken with a Leica DMS1000 microscope. For hypocotyl length measurements, seeds were germinated in white light for 8 h and then grown in vertical plates in darkness for 7 days. Hypocotyl length was measured with FIJI.

For flowering-time measurements, seeds were sown on pots, stratified for 7 days at 4°C and grown under SD (8 h light/16 h dark) or LD (16 h light/8 h dark) photoperiods at 22°C. Flowering time was recorded as the number of rosette and cauline leaves or days at bolting.

For the dormancy assay, seed lots to be compared were freshly harvested on the same day from individual plants grown in identical conditions. Seeds were sown immediately without stratification and incubated under continuous light. For the germination assay, freshly harvested seeds were sown and stratified for 3 days at 4°C. The percentage of seeds with an emerged radicle was determined at different time points.

### Stress Tolerance Assays

NaCl, LiCl, and mannitol tolerance was analyzed by transferring 4-day-old seedlings grown on vertical MS plates under LD conditions to new MS plates supplemented with or without 100 mM NaCl, 12 mM LiCl, or 300 mM mannitol and incubated vertically for 4 days. Root length was measured using FIJI. Tolerance to freezing temperatures was determined as follows: 2-week-old plants grown on plates under LD photoperiod at 20°C were transferred to 4°C for 5 days and subsequently exposed to −12°C for 6 hours. Survival rates were determined after 1 week of recovering at 20°C.

### RNA Extraction and Quantitative RT-PCR

For expression analysis of flowering-related genes, seeds were sown on soil and grown in LD. Shoot apex and the second oldest leaf from 14-day-old plants were collected. For expression analysis of auxin-related genes, seedlings were grown for 5 days under continuous light at 22°C and then transferred to 29°C for 2 h. For *PFD* expression in the presence of salt or mannitol, 7-day-old seedlings grown in continuous light were exposed to 100 mM NaCl or 300 mM mannitol in liquid media for 0, 8, or 24 h. Total RNA was extracted using Machery-Nagel kit and treated with DNase I on column (Machery-Nagel) following manufacturer’s instructions. cDNA was synthesized with the PrimeScript^™^ 1st strand cDNA Synthesis Kit (Takara), and used as a template for qPCR assays employing the SYBR^®^ Premix Ex Taq^™^ II (Takara) with primers listed in Table S1. The relative expression values were calculated using the At1g13320 (*PDF2-1*) gene as a reference, using the ΔΔCT method. All assays were performed with at least two biological replicates, each including three technical replicates.

For RNA-seq experiments, two type of samples were collected: (i) 7-day-old wild-type and *6x pfd* seedlings grown under continuous light at 22°C, and (ii) wild-type and *pfd4* seedlings grown under LD conditions for 2 weeks at 22°C. Total RNA was extracted with the RNeasy Plant Mini Kit (QIagen), and the RNA concentration and integrity (RIN) were measured in a RNA nanochip (Bioanalyzer, Agilent Technologies 2100). The preparation of the libraries and the subsequent sequencing in an Illumina NextSeq™ 500 platform was carried out by the Genomic Service of the University of Valencia.

### RNA-seq Analysis

Read trimming was performed with cutadapt. Approximately, 20 million 75 bp paired-end reads per sample were generated and > 90% reads were aligned to the TAIR10 Col-0 reference genome using HISAT2 with default parameters. htseq-count was used for read counting and DESeq2 for identifiying DEGs as those that display absolute value of log2 fold chance (logFC) > 1 and *P* adjusted value < 0.05. Raw sequences (fastq files) and differential expression gene tables used in this paper have been deposited in the Gene Expression Omnibus (GEO) database (accession no. GSE138432). Raw data from previously published RNA-seq can be found in the same database with accession no. GSE124812.

Heatmaps from RNA-seq data were performed using the http://www1.heatmapper.ca/expression/ website. Volcano plots were constructed with the EnhancedVolcano R package. GO terms were obtained from AgriGO v2 and then were filtered with REVIGO for doing scatter plots. The Integrative Genomics Viewer was used to visualize reads from alignment files.

### Microarray-based Expression Analysis

Data for *PFD* gene expression was gathered from the Bio-Analytic Resource Homepage Arabidopsis eFP Browser (http://bar.utoronto.ca/efp/cgi-bin/efpWeb.cgi). Heatmaps were generated using the Matrix2png tool (https://matrix2png.msl.ubc.ca/).

The coexpression analysis was done with the CORNET webtool (https://bioinformatics.psb.ugent.be/cornet/versions/cornet3.0/). Options used were: Pearson correlation method, correlation coefficient >0.7, retrieve top 170 genes, show pairwise correlations only. Microarray Compendium 1 TAIR10 (454 experiments with bias towards cell cycle, growth, and development) was used as source data.

### Statistical Analysis

*P* values in overlapping DEGs from two genotypes or conditions were calculated using hypergeometric tests. The rest of P values were obtained from one-way ANOVA tests followed by multiple comparison tests when more than two genotypes were compared together. t-tests were performed instead when comparing only two genotypes.

## Acknowledgements

We thank Dr. Jiri Friml (1ST, Austria) for seeds of *UBQ10::Venus-TUA6* and Dr. Karin Schumacher (University of Heidelberg, Germany) for the anti-DET3 antibody. Part of imaging analyses were carried out at NOLIMITS, an advanced imaging facility established by the University of Milan.

## Funding

This work was supported by grants from the Spanish Ministry of Economy and Competitiveness and “Agencia Española de Investigación”/FEDER/European Union (BIO2013-43184-P to D.A. and M.A.B., and BIO2016-79133-P and PID2019-109925GB-I00 to D.A.). N.B-T. and A.S.-M. were recipient of a Ministerio de Economía y Competitividad (BES-2014-068868) and EU MSCA-IF (H2020-MSCA-IF-2016-746396) fellowships, respectively.

---

**Figure S1.**
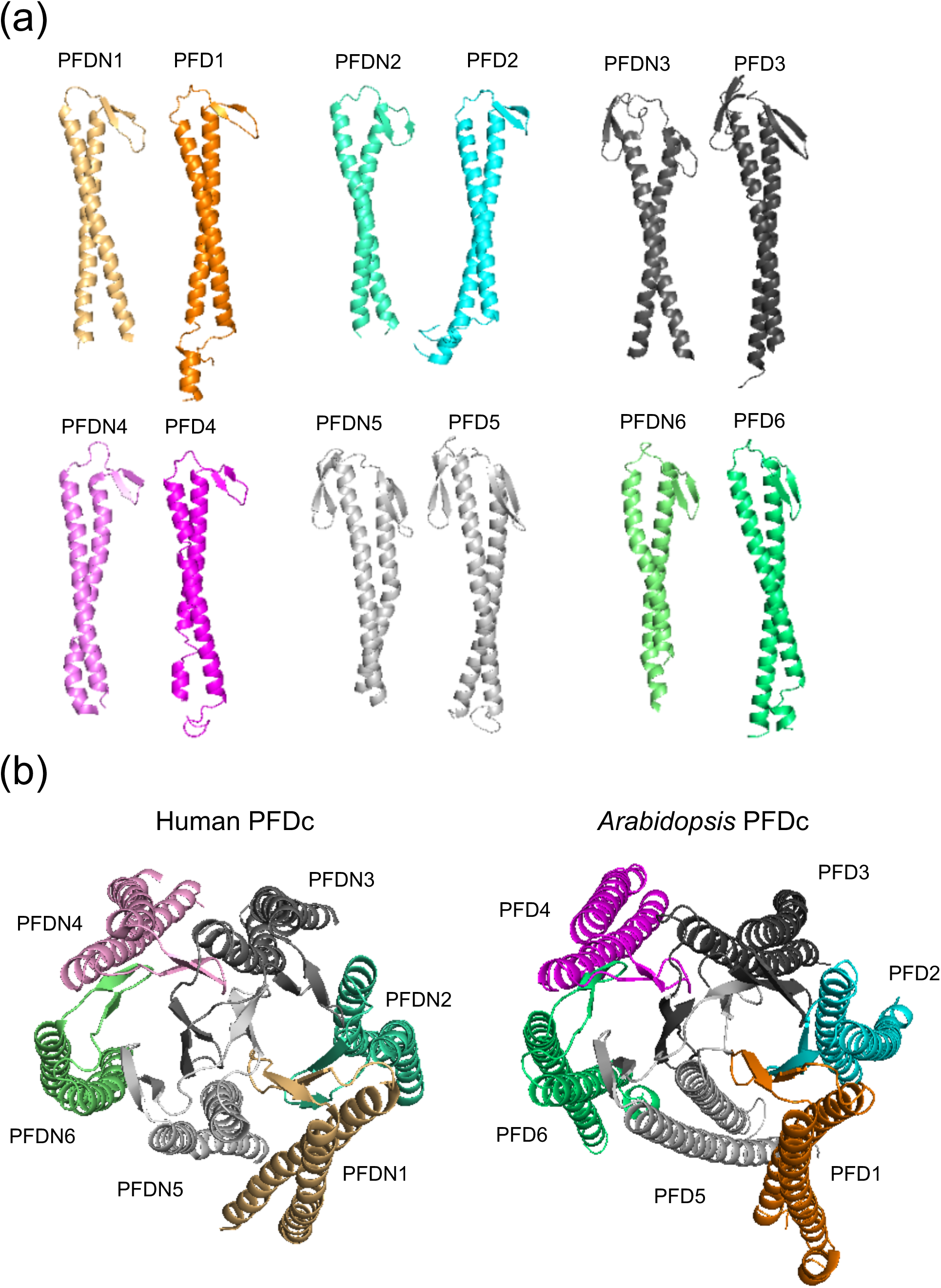
Structure of the human and *Arabidopsis* PFDc. The structure of the different *Arabidopsis* and human PFDs is shown in (a). PFDN1 to PFDN6 and PFD1 to PFD6 refer to the human and *Arabidopsis* PFDs, respectively. The 3D structure of *Arabidopsis* and human PFDc is shown in (b). The 3D structure of the *Arabidopsis* PFDc was modelled by using the human PFDc as template.

**Figure S2.**
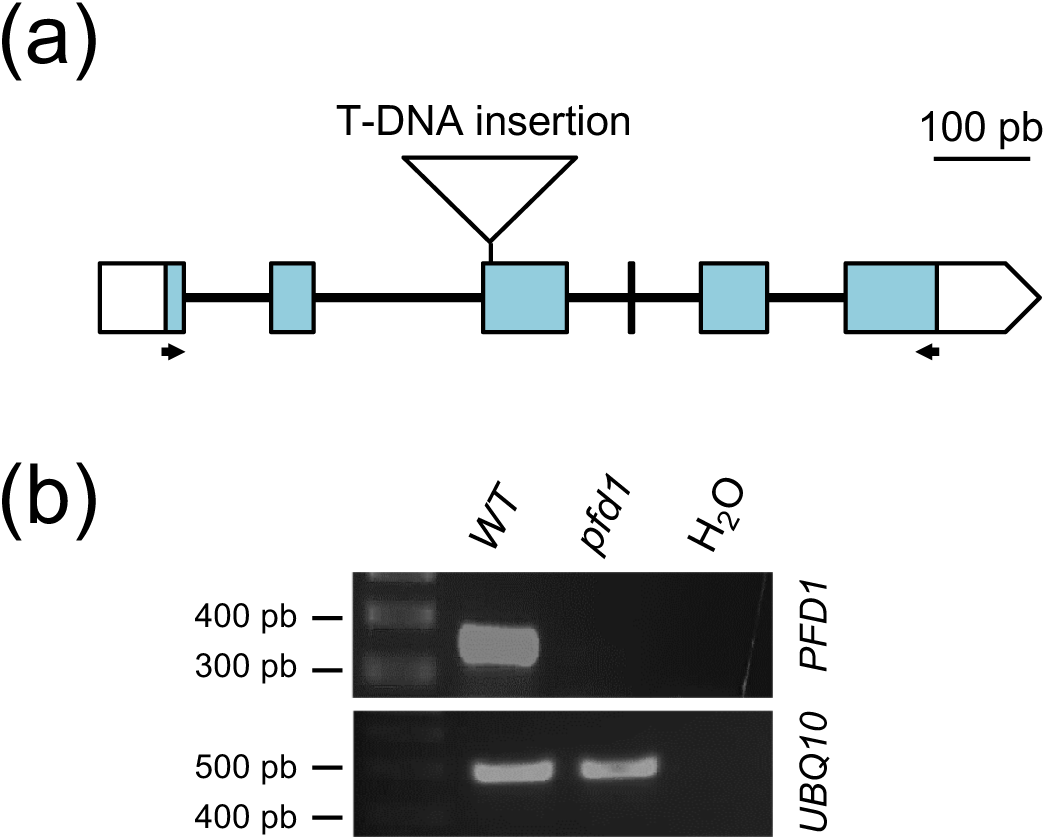
The novel *pfd1* mutant is a null allele. (a) Structure of the *PFD1* gene indicating the position of the T-DNA insertion (triangle). Boxes and lines indicate exons and introns, respectively. White boxes correspond to the 5’ and 3’ untranslated regions. Arrows indicate the position of oligonucleotides used for the semi-quantitative RT-PCR analysis of *PFD1* expression. (b) Semi-quantitative RT-PCR analysis of *PFD1* expression in wild-type (WT) and *pfd1* plants. A negative control (water) was also included. PCR products were amplified with primers indicated in Table S1.

**Figure S3.**
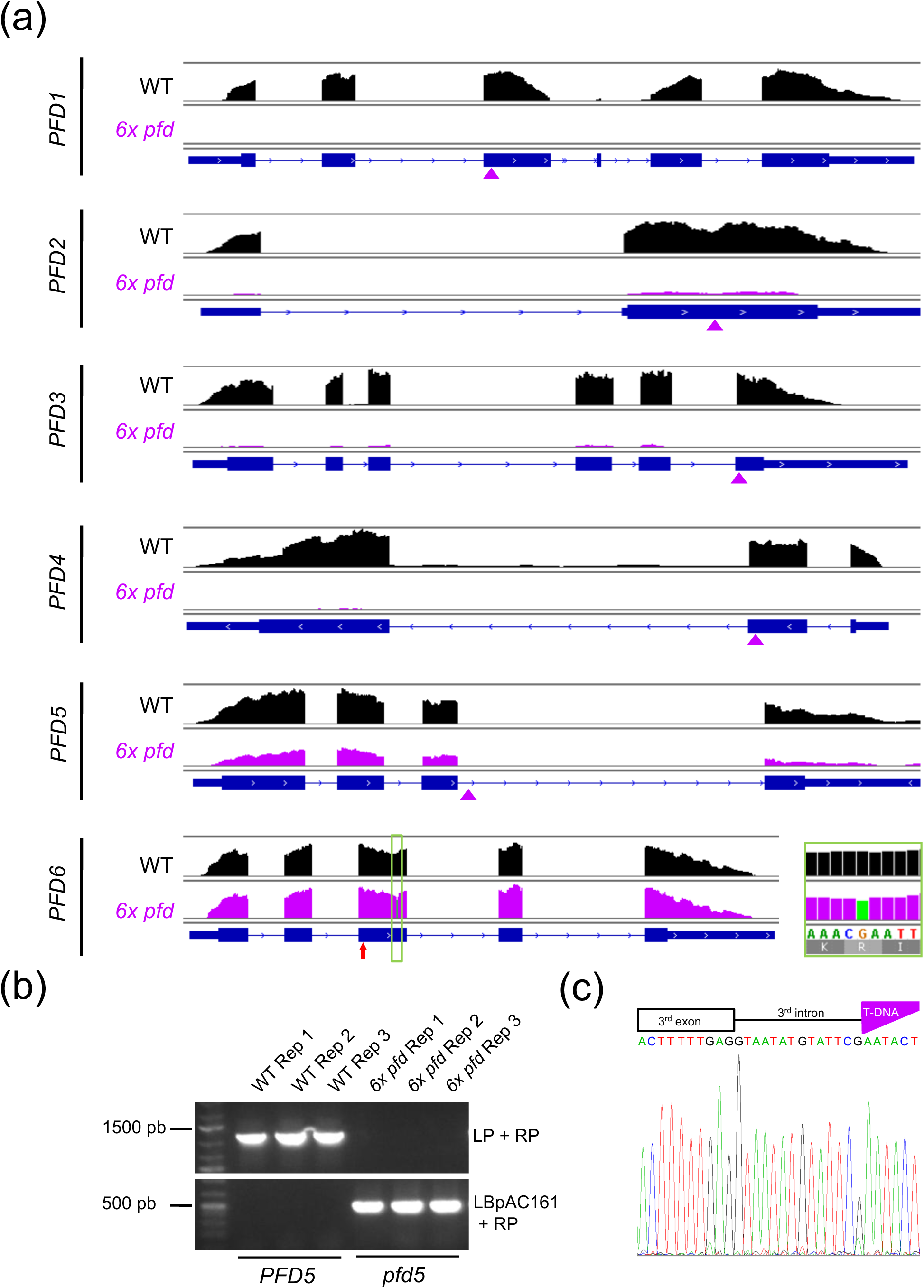
Expression of *PFD* genes in *6x pfd*. (a) IGV plots showing the RNA-seq average reads density (three replicates) in *PFD* genes in the wild type (black) and *6x pfd* (purple). Purple triangles and a red arrow indicate the position of insertions and the *pfd6-1* mutation, respectively. (b) The *6x pfd* seedlings used for RNA-seq were homozygous for the *pfd5* insertion. (c) Electrophoregram showing the position of the T-DNA insertion in *pfd5*.

**Figure S4.**
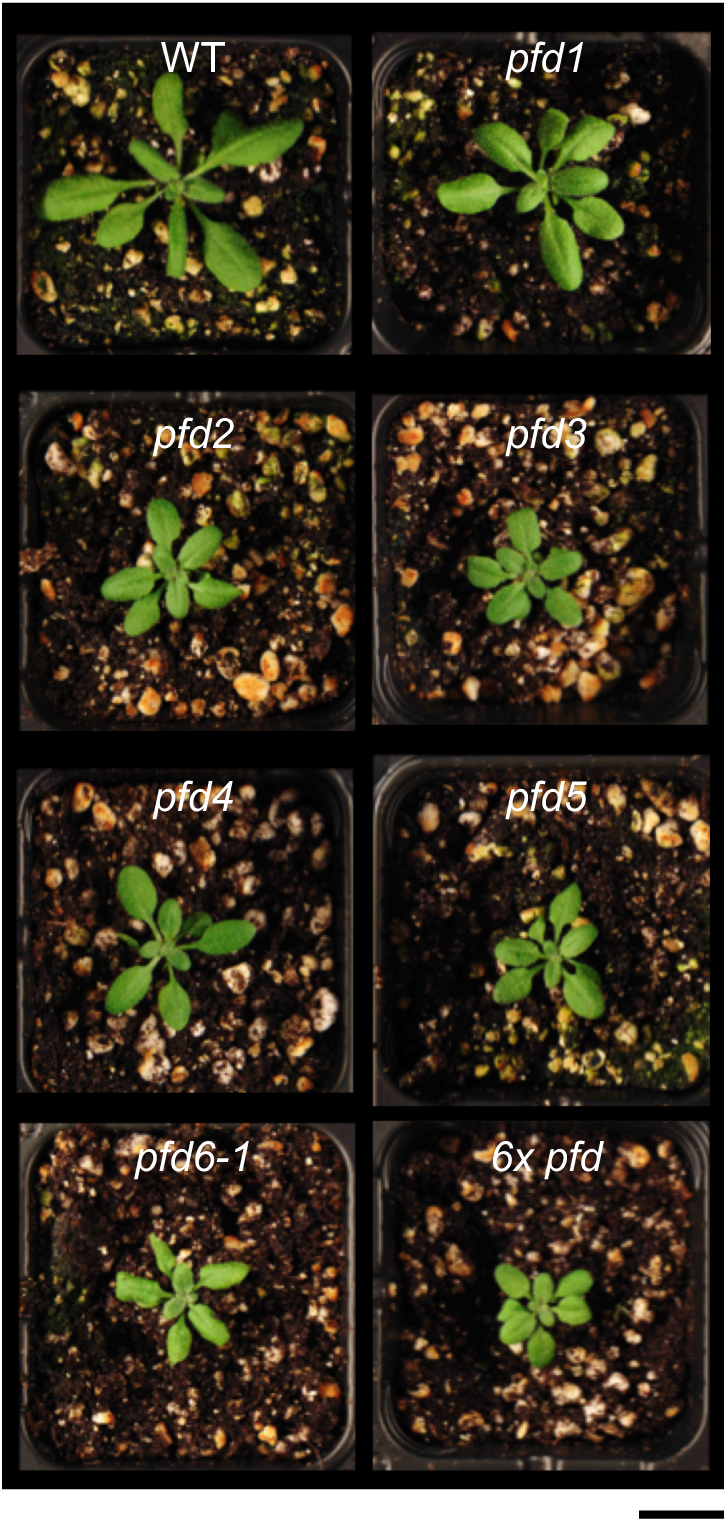
*pfd* mutant plants grown on soil. Representative images of 21-day-old plants of the indicated genotypes grown under LD conditions are shown. Scale bar = 2 cm.

**Figure S5.**
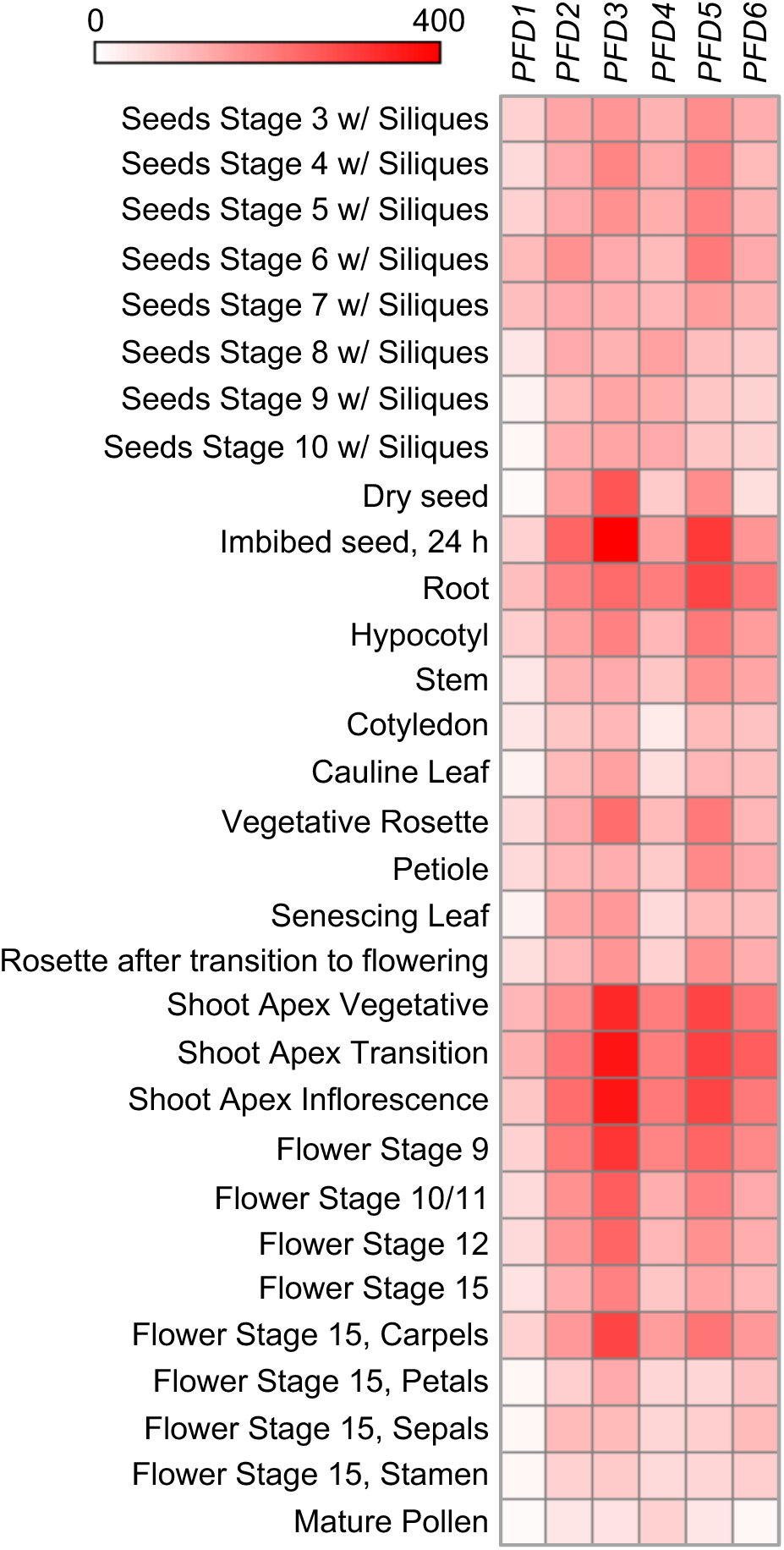
Heatmap showing the expression levels of the six *PFD* genes at different stages of *Arabidopsis* development.

**Figure S6.**
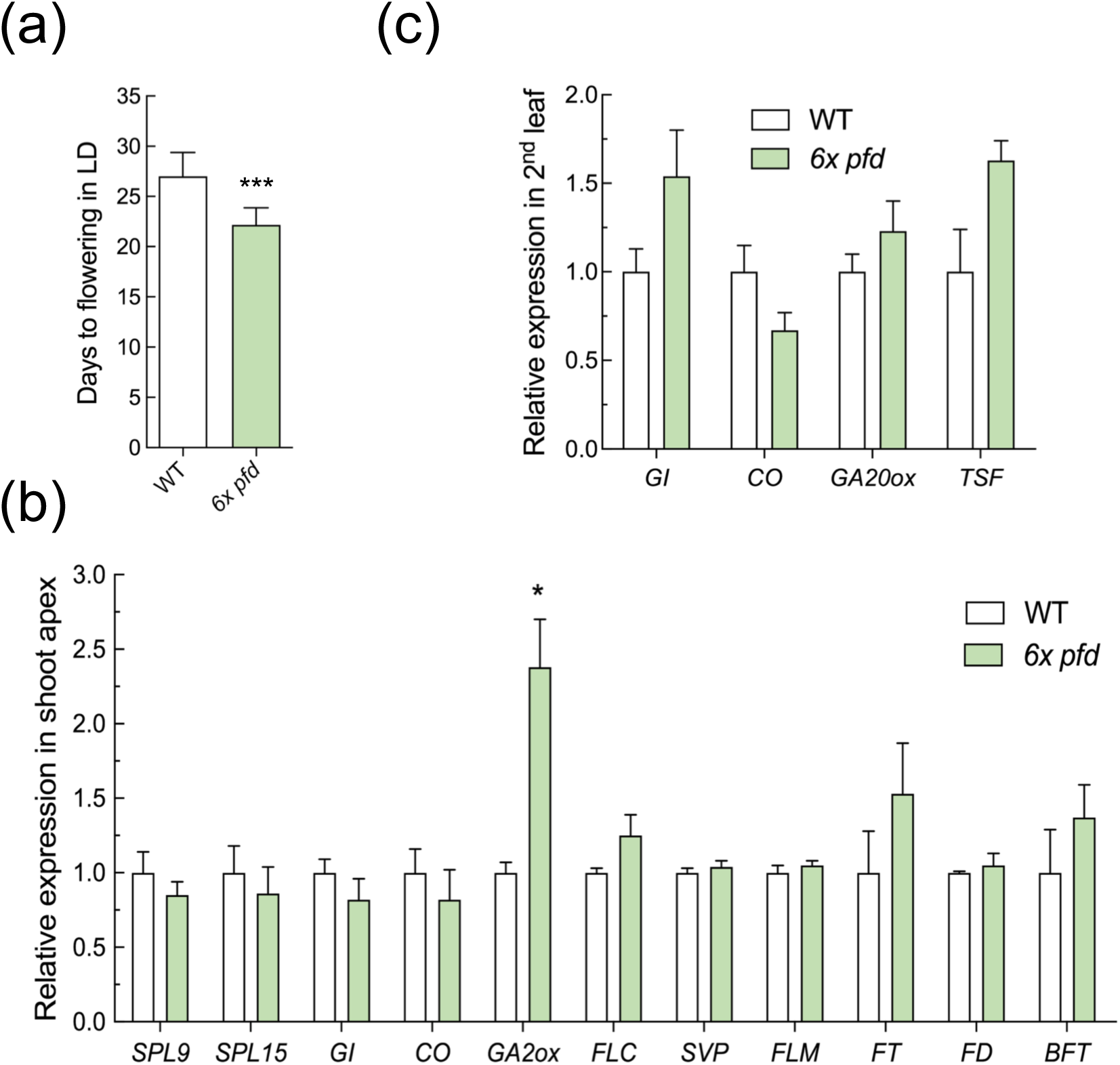
The *6x pfd* mutant flowers early. (a) Flowering time of wild-type and *6x pfd* plants (n ≥ 7) under LD, measured as the number of days for bolting. Three asterisks represent *P* < 0.001 in a t-test. (b,c) Expression analysis of flowering-time regulator genes in shoot apex (b) or the 2^nd^ oldest rosette leaf (c). Data are mean from 3 biological replicates. Error bars represent standard error from means. One asterisk represents *P* < 0.05 in t-test.

**Figure 7.**
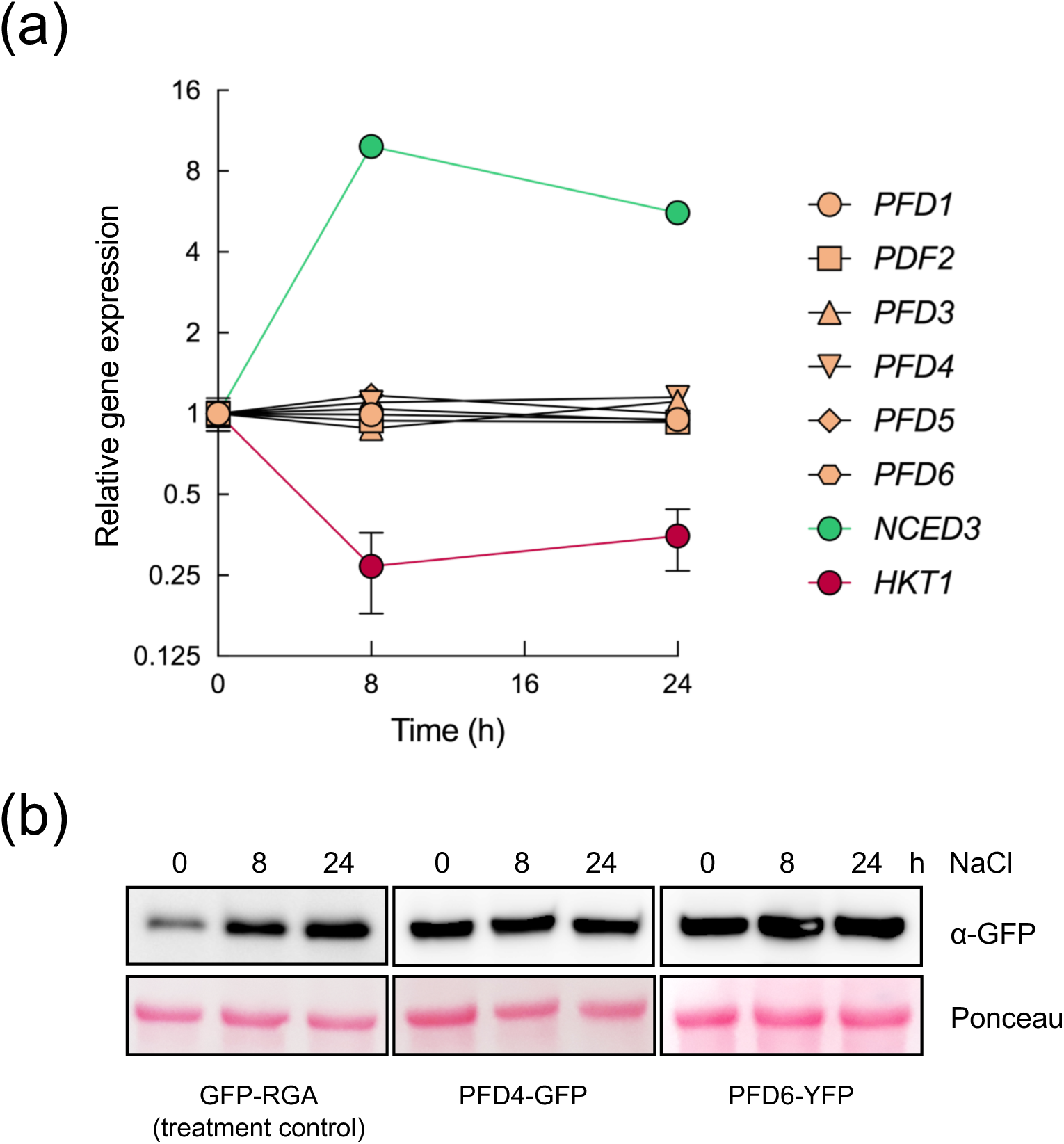
PFDs’ transcripts or proteins do not respond to salt stress. (a) Levels of *PFD1* to *PFD6* transcripts in 7-day-old seedlings grown in continuous light and exposed to 100 mM NaCl for 0, 8 or 24 hours. Values represent mean from 3 technical replicates. Error bars represent the standard deviation of these replicates. *NCED* and *HKT1* were used as control for the NaCl treatment. A second biological replicate showed equivalent results. (b) Levels of GFP-RGA, PFD4-GFP, and PFD6-YFP fusion proteins in 7-day-old seedlings grown in the same conditions than in (a).

**Figure S8.**
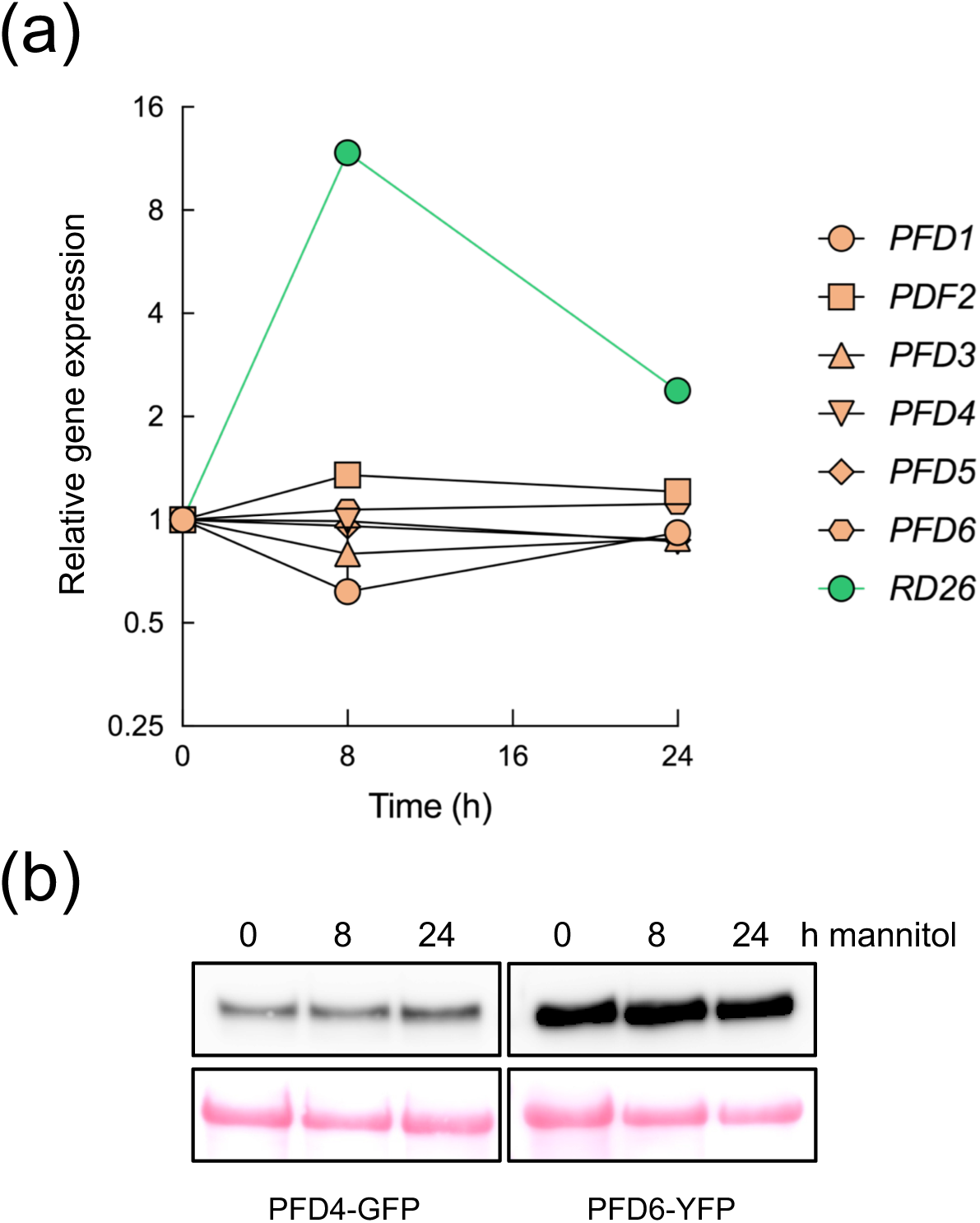
PFDs’ transcripts or proteins do not respond to osmotic stress. (a) Levels of *PFD1* to *PFD6* transcripts in 7-day-old seedlings grown in continuous light and exposed to 300 mM mannitol for 0, 8 or 24 hours. Bars represent mean from 3 technical replicates. *RD26* was used as control of the temperature treatment. A second biological sample showed equivalent results. (b) Levels of PFD4-GFP and PFD6-YFP proteins in 7-day-old seedlings grown in the same conditions than in (a).

**Figure S9.**
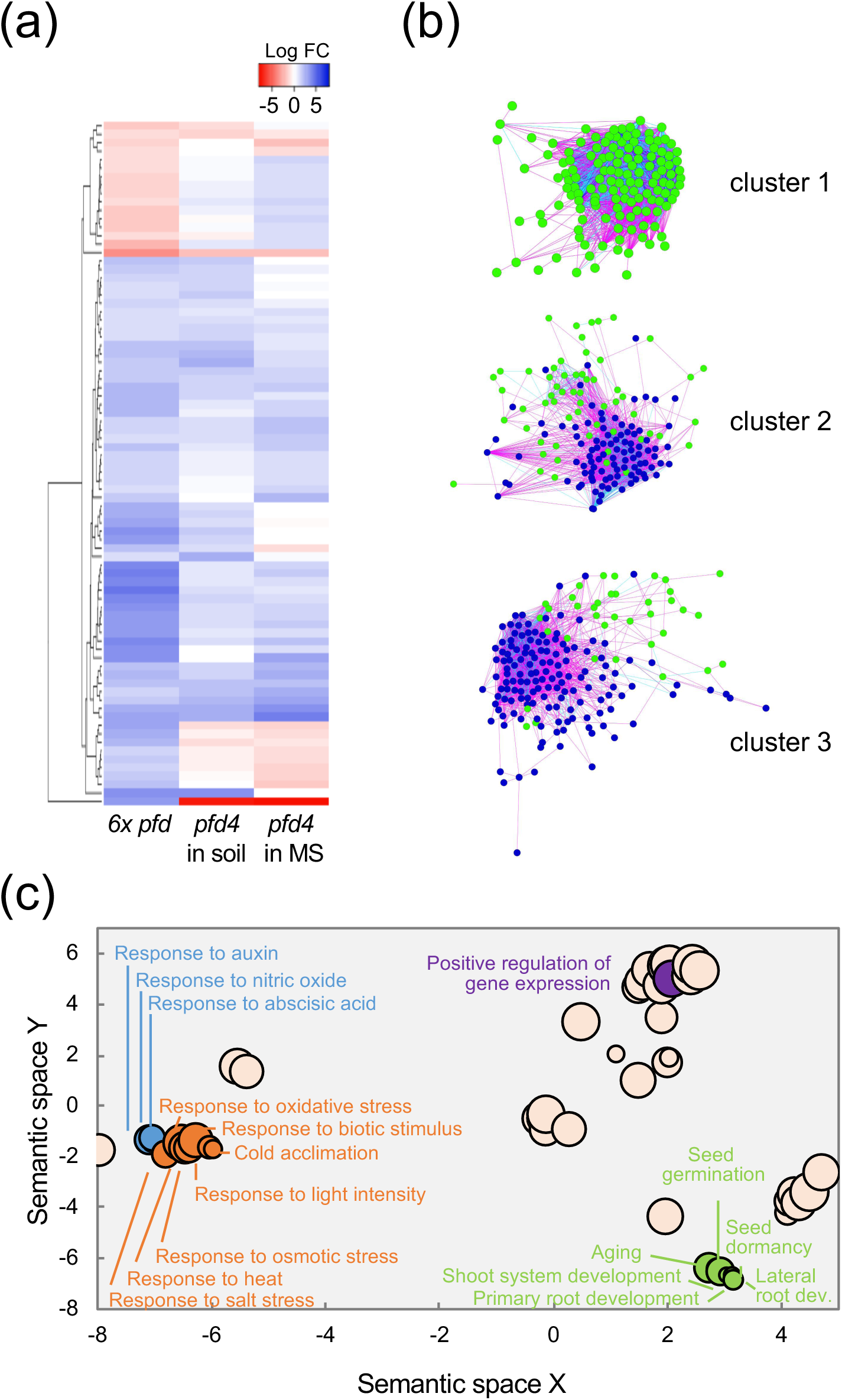
RNA-seq analysis of *6x pfd* seedlings. (a) Heatmap showing the behavior of the 81 common DEGs of *6x pfd* and *pfd4* mutants grown in control conditions. (b) A magnification of the 3 clusters shown in Figure 3c. Pink edges mean correlation 0.7-0.8, light blue edges man correlation 0.8-0.9, and dark blue edges mean correlation 0.9-1. (c) GO terms enriched in DEGs in the *6x pfd* mutant. Only some GO terms are represented. All GO terms are listed in Supplementary File 1. Bubble size is proporcional to the *P* significance of GO enrichment.

**Figure S10.**
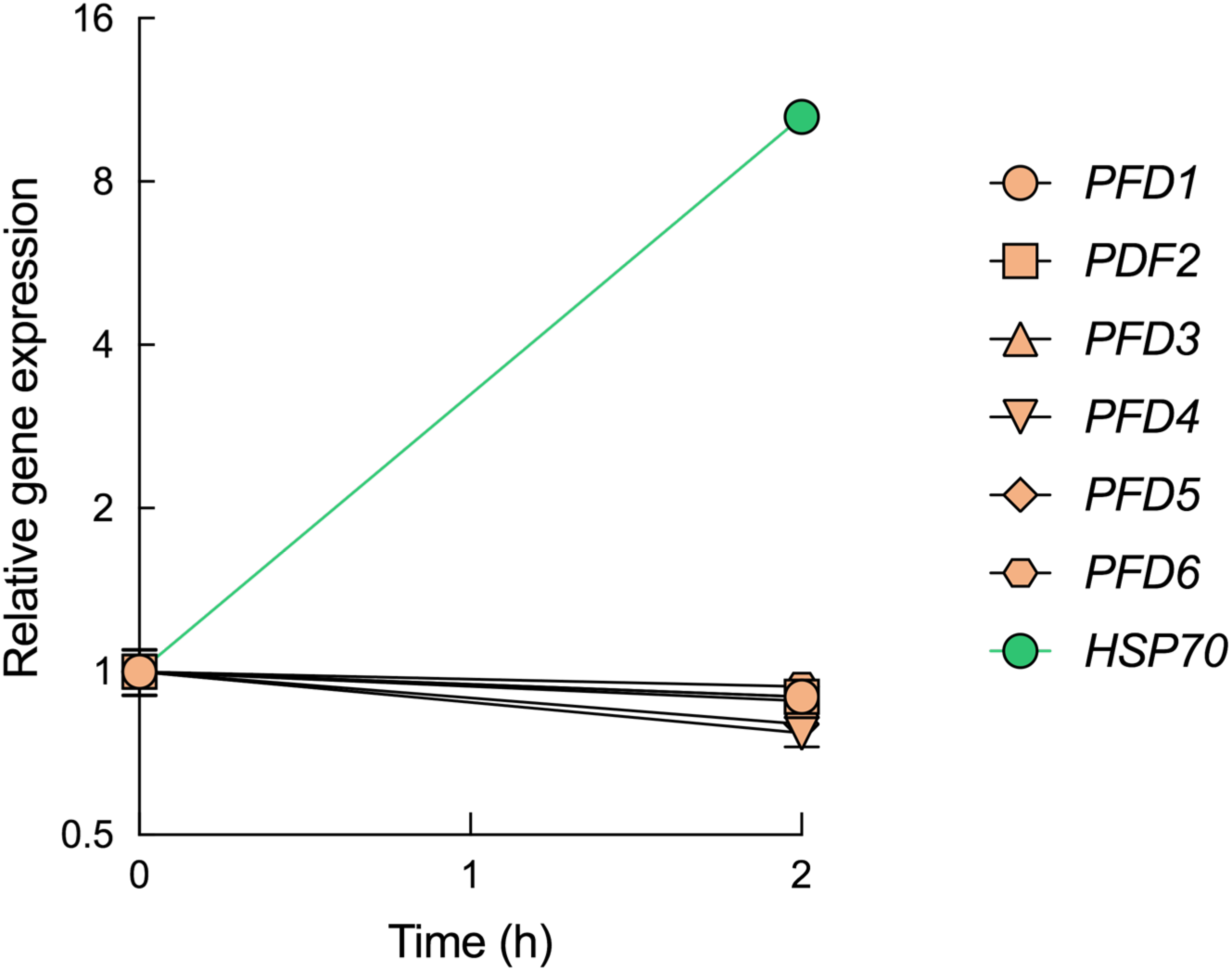
PFDs’ transcript levels do not change in responseto warm temperatures. Levels of *PFD1* to *PFD6* transcripts in 5-day-old seedlings grown in continuous light at 22°C and exposed to 29°C for 2 hours. Values represent mean from 3 technical replicates. Error bars represent the standard deviation of these replicates. A second biological sample showed equivalent results. *HSP70* was used as control for the temperature treatment.

**Figure S11.**
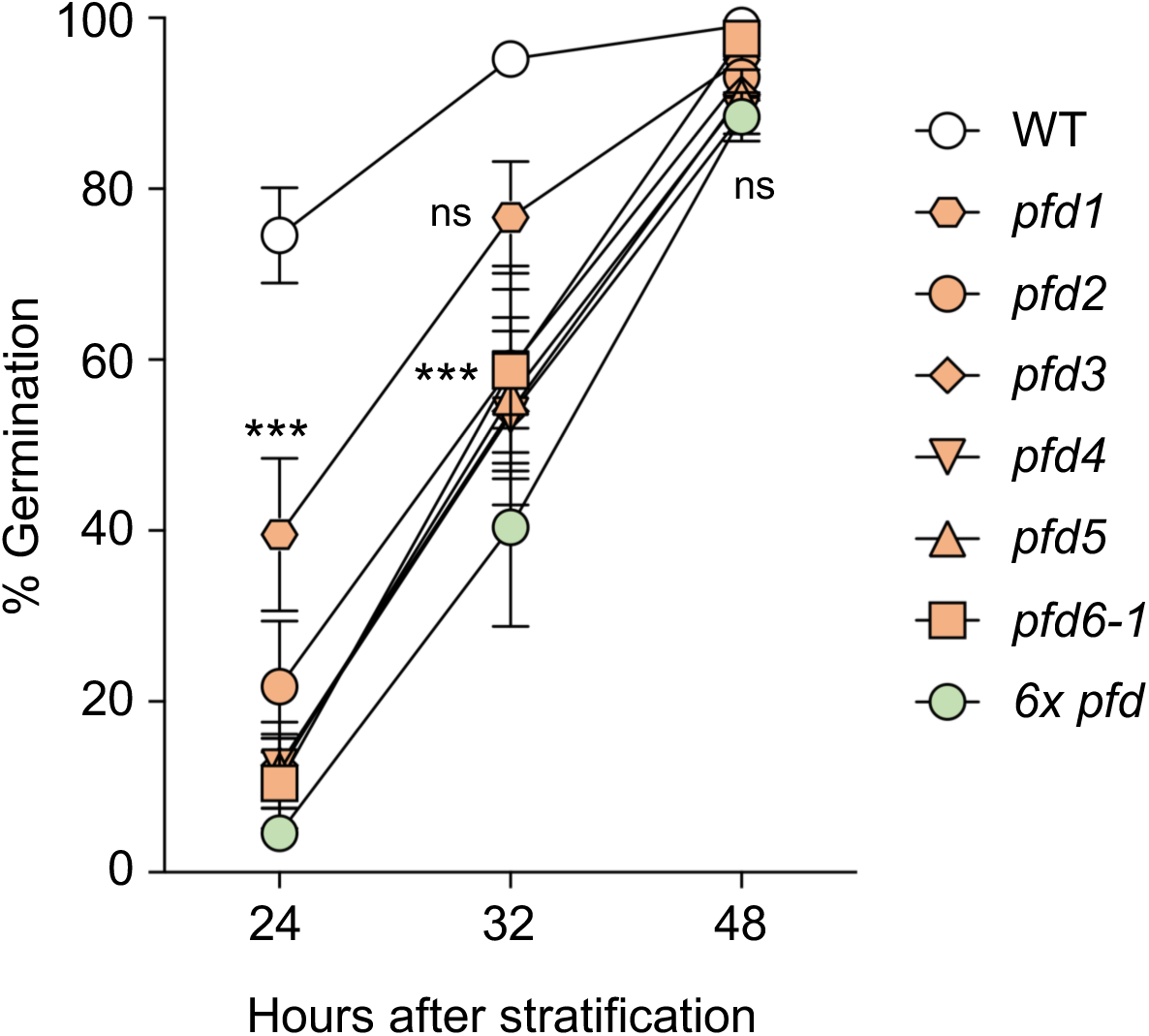
The PFDc contributes to seed germination. Germination rates of stratified seeds. Error bars represent standard error of mean from seven biological replicates (each one including at least 29 seeds). Three asterisks represent *P* < 0.001 in Bonferroni tests after ANOVA tests; when comparing with the wild type, all *pfd* mutants show this significant difference at 24 h, while at 32 h the *pfd1* shows no significant differences. *ns*, no significant differences.

**Figure S12.**
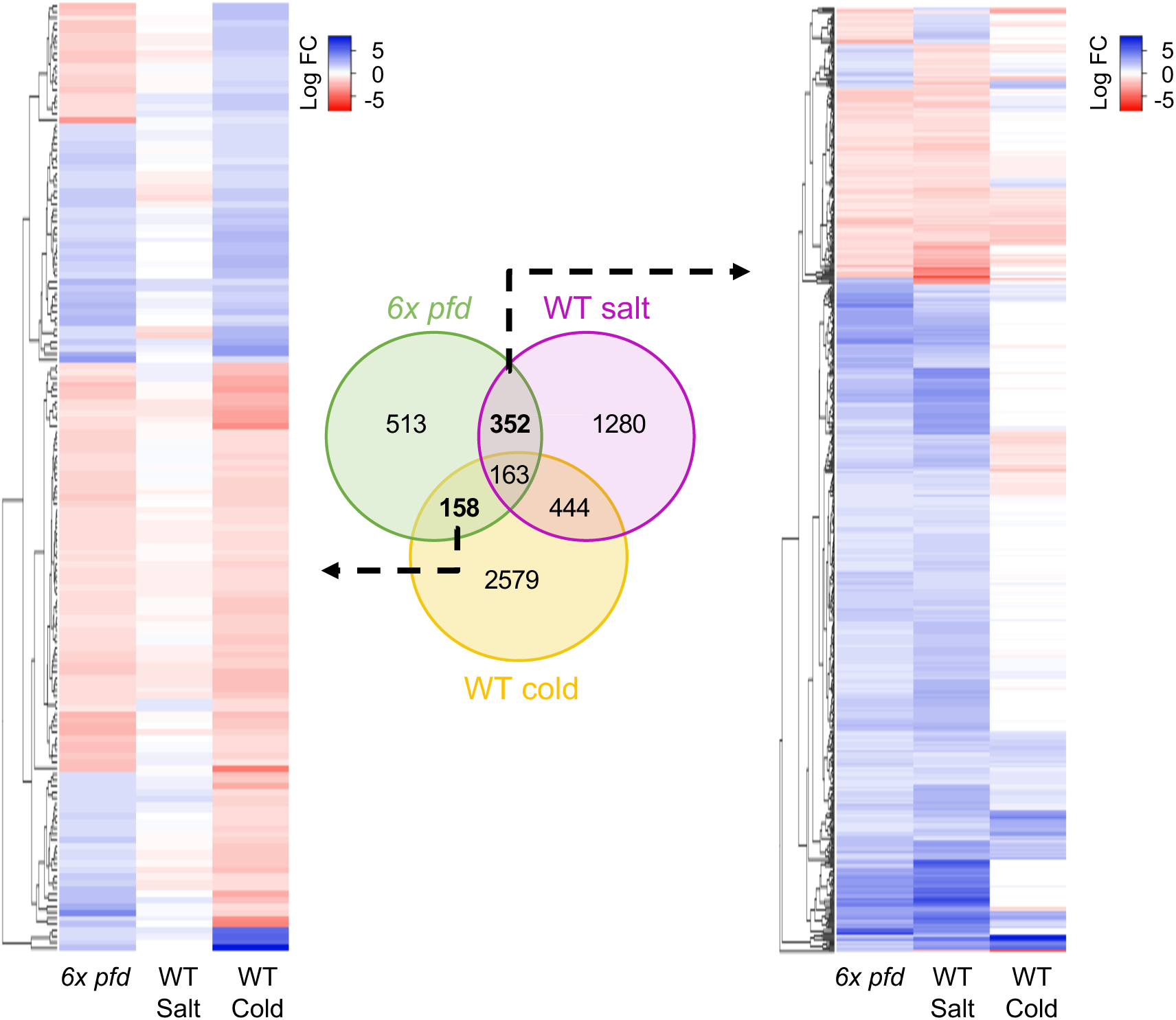
Constitutive stress signature in *6x pfd* seedlings. The Venn diagram shows the overlap between DEGs. Heatmaps show the behavior of DEGs.

**Figure S13.**
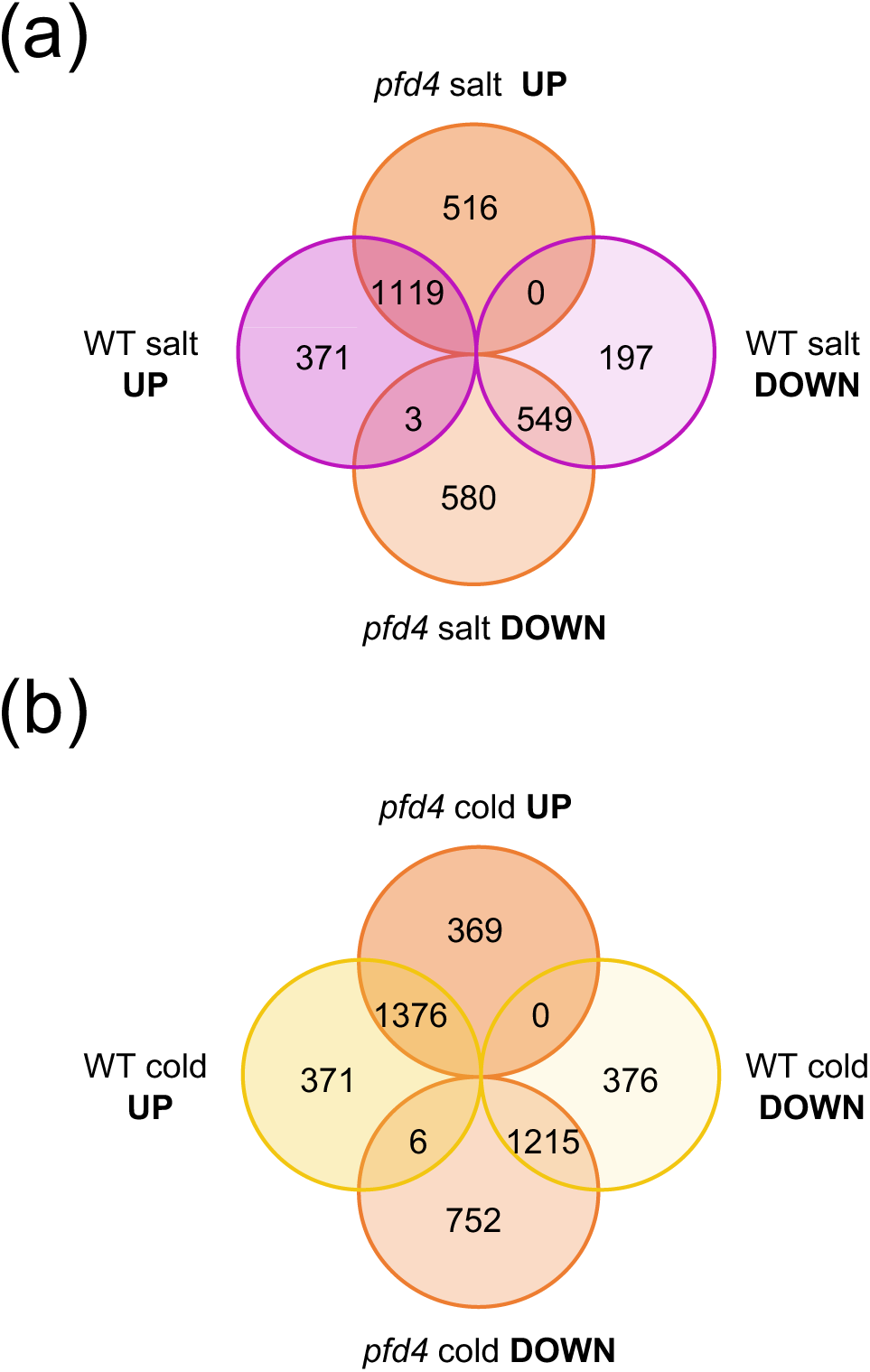
PFD4 is involved in the regulation of the gene expression in response to cold and salt stresses. Venn diagrams showing the overlaps between DEGs in the wild type and *pfd4* seedlings in response to high salt (a) or low temperature (b).

## Notes

### Competing Interest Statement

The authors have declared no competing interest.

