## Supplemental Table for "A genetic approach to dissect the role of prefoldins in *Arabidopsis*"

| **Table S1.** Oligonucleotides used in this work | | | |
| --- | --- | --- | --- |
| Purpose | Name | Oligonucleotide sequence (5’ 🡪 3’) | |
|  |  | Forward primers | Reverse primers |
| Genotyping | pfd1-LP/RP^a^ | AATCCCAAAAGCTCTCCAAAC | ATCAAGATCTGCAGACATGGC |
|  | pfd2-LP/RP^b^ | AACGATTGGAGAAGCATTGG | ACAATTGTGTTTTTCGGTTCG |
|  | pfd3-LP/RP^a^ | GCTAAAGCCAGCCTGGAGGTTCTTG | cggaaggcaagcgatgataacaatg |
|  | pfd4-LP/RP^c^ | ATCCCCCTAATGAAACGATTG | TAACCAATTCCAAGCCAAGTG |
|  | pfd5-LP/RP^a^ | tagtctcaatagcaaaaagcac | TCAAGATAGTCTCAACAACATCCG |
|  | pfd6-1-TaqI-F/R^d^ | GAAGCTAATGCAAATGTTCGTAATC | ACTTATGAACAGCAATCACTCACT |
|  | Venus-TUA6-F/R | ACCTACGGCAAGCTGACCC | AGCCTGACCAATGTGGATCG |
|  | LBpAC161^a^ | ATATTGACCATCATACTCATTGC |  |
|  | LB4^b^ | TGATCCATGTAGATTTCCCGGAC |  |
|  | LBb1.3^c^ | ATTTTGCCGATTTCGGAAC |  |
| Semi  q-PCR | PFD1 -F/R | ATGGCGGACGAAGCAACCAG | CATAGACATGGACATAATCTGTTG |
|  | UBQ10-F/R | GATCTTGCCGGAAAACAATTGGAGGATGGT | CGACTTGTCATTAGAAAGAAAGAGATAACAGG |
| qPCR | qPFD1-F/R | CGAAAGACCCTTCTCTAGCACGTCG | CTCCATAAAAGCAGCTCTGGTTGC |
|  | qPFD2-F/R | GAAGCGGTGGCCTAAGAGAACC | CTCACTTGCATCTCGAGGTCGG |
|  | qPFD3-F/R | CCCCTACTGCTATTGCTGTTGCAG | CAACCTGATTGAGGCAAAAGGTGTCC |
|  | qPFD4-F/R | ACAACTCTCTCTCACACAGCAACGTTC | GATCCACTCTTACTCCCTTGCTGC |
|  | qPFD5-F/R | GAAAAGCGTGGCAGATGAAGCTGG | TCACGACGTGGTTGCAGCTG |
|  | qPFD6-F/R | CCAACTCGGTGAGAATGAGCTCG | CCAAATCCTGCTTCACTAACACCGG |
|  | qPDF2-1-F/R | TAACGTGGCCAAAATGATGC | GTTCTCCACAACCGCTTGGT |
|  | qNCED3-F/R | CGGTGGTTTACGACAAGAACAA | CAGAAGCAATCTGGAGCATCAA |
|  | qHKT1-F/R | GAAAGGCAAAATCTACAACGTG | CCTGCAAACCCATAACTCG |
|  | qHSP70-F/R | GAAGTACAAGGCTGAGGATGAAGAAC | CTTCTCGTCCTTGATCGTGTTCC |
|  | qRD26-F/R | CATCGTCTTCTTCATCACAG | TAAGACCTGCCAAGCTAG |
|  | qYUC8-F/R | AAACGCTCAAGGGGTTCTCTTCG | CACGCACAACACCCTTTGATTCG |
|  | qIAA19-F/R | GGTGACAACTGCGAATACGTTACCA | CCCGGTAGCATCCGATCTTTTCA |
|  | qIAA29-F/R | AAACAGCGTTTGTTTGCCTTGAATG | TGGCCATCCAACAACTTCGCTAT |
|  | qSAUR19-F/R | CTTCAAGAGCTTCATAATAATTCAAACTT | GAAGGAAAAAATGTTGGATCATCTT |
|  | qSAUR23-F/R | ATTCAAACTTTCAGACAAAAGAAATGG | ACAAGGAAACAACTCTATCTCTAACT |
|  | qSPL9-F/R | GGAATTTGACCTAGAGAAAAGGAGTT | GCATCACCATTTTCGTAAAGCGAAG |
|  | qSPL15-F/R | TGAATGTTTTATCACATGGAAGCTC | TCATCGAGTCGAAACCAGAAGATG |
|  | qGI-F/R | AGCAGTGGTCGACGGTTTATC | ATGGGTATGGAGCTTTGGTTC |
|  | qCO-F/R | CACTACAACGACAATGGTTCC | GGTCAGGTTGTTGCTCTACTG |
|  | qGA20ox2-F/R | CGAGCAGTTTGGGAAGGTGTATC | CCTAAACTTAAGCCCAGAAGCTCC |
|  | qGA2ox2-F/R | GGTTCCGGTTCTCACTTCCC | GGATCGGCTAGGTTGACGAC |
|  | qFLC-F/R | CTTGTGGGATCAAATGTCAAAAATGTG | CATCTCAGCTTCTGCTCCCACATGATG |
|  | qSVP-F/R | CAAGGACTTGACATTGAAGAGCTTCA | CTGATCTCACTCATAATCTTGTCAC |
|  | qFLM-β-F/R | CATGCTGATGAACTTAGAGCCTTAGATC | CAGCAACGTATTCTTTCCCAT |
|  | qFT-F/R | CCCTGCTACAACTGGAACAAC | CACCCTGGTGCATACACTG |
|  | qTSF-F/R | TGCCACCACTGGAAATGCC | CGTTTGTCTTCCGAGTTGCC |
|  | qFD-F/R | GCTCACTTGCAGGCAGAAAA | CCTTTTCTCTTTCCGGGTCT |
|  | qBFT-F/R | CGCCGGAAACTAGAGAGTGT | GTTGGGCGTTGAAGTAAACA |
|  | qSOC1-F/R | AAACGAGAAGCTCTCTGAAAAG | AAGAACAAGGTAACCCAATGAAC |
|  | qLFY-F/R | ACGCCGTCATTTGCTACTCT | CTTTCTCCGTCTCTGCTGCT |
|  | qAP1-F/R | GAAGGCCATACAGGAGCAAA | ACTGCTCCTGTTGAGCCCTA |

^a,b,c^RP + the corresponding LB primer were used for genotyping insertions. ^d^The *pfd6-1* point mutation was genotyped by dCAPS using the indicated primers and the TaqI restriction enzyme.
